# The RND family efflux pump FemT contributes to lipid homeostasis in *Staphylococcus aureus”*

**DOI:** 10.64898/2026.08.10.744025

**Authors:** Akanksha Thukral, Camryn Bonn Dunbar, Jamie Halucha, James Schneider, Tyler R. Pereira, John K. McCormick, David E. Heinrichs, Martin J. McGavin

## Abstract

The RND efflux pump FemT encoded by SAUSA300_2213 of *Staphylococcus aureus* USA300 is co-transcribed with *femX* which has an essential role in synthesizing the Lipid II precursor of peptidoglycan. Anticipating that this arrangement reflects a critical accessory role for *femT*, we constructed USA300Δ*femT* to assess its function. Although growth of USA300Δ*femT* in tryptic soy broth (TSB) was not impaired, transcriptomic data revealed a mild cellular stress response, accompanied by reduced expression of *ohyA* and *crt* genes involved in fatty acid metabolism and carotenoid lipid synthesis respectively. Accordingly, USA300Δ*femT* exhibited impaired growth on exposure to saturated and unsaturated fatty acids, and exposure to subinhibitory 50 µM palmitic acid promoted accumulation of reactive oxygen species, reduced respiratory activity, and altered membrane function and morphology. The transcriptome of cells grown under this condition revealed strongly attenuated expression of *ohyA* and *crt*, and several genes required for oxidative and anaerobic respiration, concomitant with strongly enhanced expression of several stress response pathways. Cellular metabolites were also profoundly altered. Finally, lipidomic analysis of USA300Δ*femT* exposed to oleic acid revealed increased incorporation of oleic acid into phosphatidylglycerol, accompanied by a significant reduction in undecaprenol C55 lipid carrier, and respiratory quinones MK-7 and MK-8. Our data are consistent with a role for FemT in maintaining cellular lipid homeostasis by promoting efflux of isoprenoid and carotenoid lipids that are prone to oxidative damage, including C55 and menaquinones that undergo cyclic reactions in peptidoglycan synthesis and electron transport.

**IMPORTANCE:** The FemT efflux pump of *S. aureus* is co-expressed in an operon with *femX* encoding an essential enzyme needed to complete the synthesis of peptidoglycan precursor Lipid II. Although this alluded to a specific role for FemT in supporting peptidoglycan synthesis, our data are instead consistent with a general role in efflux of cellular isoprenoids and carotenoid lipids that are susceptible to oxidation during routine cellular functions. Consequently, *S. aureus* became strongly dependent on FemT function when exogenous host-derived fatty acids were being actively metabolized. This represents a significant advance in our understanding of the role of an RND efflux pump in supporting routine growth-related functions of *S. aureus* and exposes a function that could be targeted to impair *S. aureus* growth on exposure to host-derived fatty acids.

## INTRODUCTION

*Staphylococcus aureus* is a major leading cause of bacterial infection related mortalities worldwide, second only to *Mycobacterium tuberculosis* in consistently exceeding one million attributed deaths annually (Piewngam and Otto, 2024). Multidrug resistance is increasingly reported in *S. aureus* (Thacharodi *et al*., 2025) and in 2024 methicillin resistant *S. aureus* (MRSA) was recognized by the World Health Organization as a public health threat, especially in health care settings (Sati *et al*., 2025). In pursuit of tackling infections caused by multidrug resistant pathogens, development of newer antibiotics including inhibitors to multidrug transporters have gained momentum, as these transporters can expel a wide range of antimicrobial agents, thereby reducing drug efficacy (Santajit and Indrawattana, 2016; Alenazy, 2022). This is especially true for resistance nodulation division (RND) superfamily efflux pumps in Gram-negative pathogens, which promote efflux of most clinically relevant antibiotics (Blair et al., 2015; Hayashi et al., 2015). With a focus on antimicrobial resistance, the mechanistic and structural details on substrate efflux through proton motive force are well known (Yamaguchi, Nakashima and Sakurai, 2015; Kavanaugh et al., 2024), but there has been less emphasis on function in microbial physiology, including identification of primary physiologic or environmental substrates. Nevertheless, a general theme has emerged that RND pumps in Gram-negative bacteria efflux diverse substrates, allowing bacteria to thrive in different environmental conditions (Tsukagoshi and Aono, 2000; Piddock, 2006)

RND pumps are less well studied in Gram-positive bacteria, perhaps due to their yet being no affiliation with resistance to antibiotics in major pathogens that include *S. aureus*, *Streptococci*, and *Enterococci*, which instead have well documented resistance mechanisms attributed to the major facilitator, ABC, and MATE family of transporters (Kaatz, McAleese and Seo, 2005; Truong-Bolduc *et al*., 2005; DeMarco *et al*., 2007). The apparent lack of emergence of RND-mediated resistance mechanisms in Gram-positive bacteria may be attributed to differences in cell envelope architecture, and the consequent unique relationship to RND efflux pump function in Gram-negative bacteria. Notably, structural studies on Gram-negative RND efflux pumps reveal that in addition to structural features in the periplasmic domain required for efflux, there are also conserved features required for complex formation with a periplasmic adaptor protein (ie; AcrA) and outer membrane protein (ie; TolC) as noted for the prototypic member of the RND efflux pump AcrB (Colclough *et al*., 2020; Athar *et al*., 2023), where these interactions ensure a contiguous channel between the efflux domain and outer membrane.

In our own research on RND efflux pump function in *S. aureus*, we and others identified a role for FarE in promoting efflux of unsaturated fatty acids and other lipid products that are toxic to the bacteria (Kenny *et al*., 2009a; Alnaseri *et al*., 2015; Huang *et al*., 2022). Consistent with Gram-positive RND efflux pumps functioning in the context of a different cell envelope architecture, FarE most closely resembles MmpL3 in *Mycobacterium tuberculosis* which promotes export of trehalose dimycolate lipid, and structural studies confirm that relative to Gram-negative RND efflux pumps, MmpL3 has a smaller extra-cytoplasmic domain lacking the features required for protein interactions in Gram-negative RND pumps (Alnaseri et al., 2015; C. C. Su et al., 2019; Adams et al., 2021). However, we also note research describing an RND efflux pump SwrC (SrfP/YerP) of *Bacillus subtilis*, which confers resistance to and export of the lipopeptide surfactin, and was noted to have high sequence similarity (46% identity) to an efflux pump in *S. aureus* (AF105976) (Tsuge, Ohata and Shoda, 2001). This efflux pump was previously characterized in *S. aureus* in context of its co-expression with *femX* encoding an essential glycyltransferase needed for synthesis of the Lipid II precursor of peptidoglycan (Quiblier et al., 2013).

Although this co-expression with *femX* alluded to a potential role in cell wall biogenesis as noted for MmpL3 in *M. tuberculosis* (C. C. Su *et al*., 2019; Su *et al*., 2021), inactivation of the gene in *S. aureus* Newman did not cause any apparent defect in growth or resistance to a range of antimicrobial agent (Quiblier et al., 2013). Given the role of SwrC in resistance to a surfactin lipopeptide, together with its reported homology to an efflux pump in *S. aureus* that is co-expressed with *femX* involved in synthesis of Lipid II, we undertook a more detailed examination of its function in *S. aureus*. We designate this gene as *femT* to denote its function as a transporter associated with FemX. Herein, we describe a phenotype for a *femT* deficient mutant in *S. aureus* strain USA300 that is consistent with a role in maintaining membrane homeostasis, and surprisingly FemT has structural features as predicted by AlphaFold that closely resemble RND efflux pumps in Gram negative bacteria, thereby blurring the association of structural details in RND efflux pumps with different cell envelope architectures in Gram-negative and Gram-positive bacteria.

## RESULTS

### Bioinformatic analysis of FemT and its paralog in *Bacillus subtilis*

The RND efflux pump encoded by SA2056 in strain Newman is co-transcribed with *femX* (Quiblier et al., 2013). *femX* encodes an essential glycyl transferase that initiates synthesis of the pentaglycine cross bridge required for completion of the Lipid II precursor of peptidoglycan which is assembled on a lipid carrier undecaprenol phosphate (C55P). In *S. aureus* USA300 FPR3757 we designated this gene (SAUSA300_2213) *femT*, due to it being an RND transporter associated with *femX.* For structural comparisons, we submitted the Alphafold structure of FemT (AF-A0A0H2XER4-F1) to DALI and searched for related proteins of known structure. This revealed that FemT has structural features inherent to RND efflux pumps in Gram-negative bacteria, with the closest match being the multidrug efflux pump AdeB of *Acinetobacter baumanii* (Figure S1 and Table S1). It was also previously noted that SwrC (SrfP/YerP) of *Bacillus subtilis* had high sequence similarity (46% identity) with an efflux pump in *S. aureus* (AF105976) corresponding to what we have defined as FemT (Tsuge, Ohata and Shoda, 2001; Kearns *et al*., 2004). FemT and SwrC also share a high degree of structural relatedness, matching that of AdeB-FemT (Fig. S1 and Table S1), whereas FarE of *S. aureus* which has an established function in lipid efflux is distinct from FemT and closely resembles MmpL3 of *M. tuberculosis* which promotes transport of trehalose monomycolates (Alnaseri *et al*., 2015; C.-C. Su *et al*., 2019). Of additional interest, *swrC* in *B. subtilis* is followed by *dgkB* (Jerga *et al*., 2007) encoding an essential diacylglycerol kinase required for lipoteichoic acid synthesis, maintaining the co-association of an RND efflux pump with an essential cell envelope synthetic function as noted with *femXT*.

### FemT is expressed throughout growth and does not respond to exogenous fatty acids

A Western blot assay for detection of FemT in lysates from *S. aureus* cells taken at different time points revealed a consistent level of production throughout growth (Fig. 1A), consistent with its expression being dependent on *femX* which is required for peptidoglycan synthesis. To confirm the dependency of *femT* expression on its co-association with *femX*, we constructed *femT::lux* and *femX::lux* reporters, where luciferase expression is under control of any potential promoter elements within *femXT* intergenic segment, or upstream of *femX,* respectively. Consistent with the expected operon structure, *femX::lux* showed consistent expression from 1-8h of growth, whereas *femT::lux* was inactive (Fig. 1B). A Western blot for detection of FemT in cultures grown in TSB, or TSB supplemented with exogenous saturated palmitic acid or unsaturated linoleic acid did not reveal any obvious differences in FemT production (Fig. 1C). These data are consistent with FemT being co-expressed in association with the essential growth-related function provided by FemX, and expression is not modulated in response to exogenous fatty acids.

**Figure 1.**
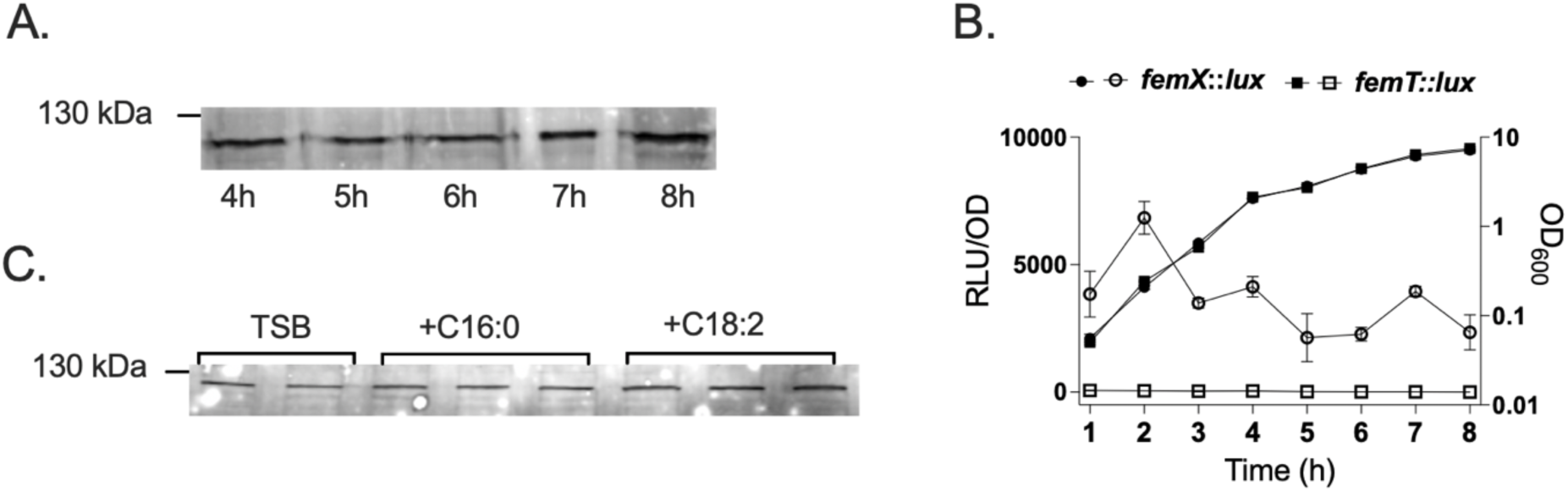
Western blot analysis of FemT expression during growth in TSB (A). Luciferase reporter assay measuring *femX* and *femT* promoter activity throughout growth, expressed as relative luminescence (RLU/OD_600_) together with the corresponding growth curves (OD_600_) (B). Western blot analysis of FemT expression following growth in TSB alone or TSB supplemented with palmitic acid (C16:0) or linoleic acid (C18:2) (C). For panel (A), USA300 was cultured in TSB alone, and samples were withdrawn at the indicated time points for preparation of whole cell lysates. 20 µg of protein was subjected to SDS-PAGE and Western blot for detection of FemT with rabbit polyclonal antiserum. (B), USA300 harboring either the *femX::lux* or *femT::lux* intergenic segment reporter gene constructs were grown in TSB in flask culture. At the indicated time points, culture optical density (OD_600_) was measured, and aliquots (4 × 200 µL technical replicates) were transferred to black 96-well microtiter plates for quantification of luciferase activity. Luciferase activity was normalized to cell density (RLU/OD_600_). Open symbols represent normalized luciferase activity (RLU/OD_600_), whereas closed symbols represent the corresponding culture growth (OD_600_). Data are presented as the mean ± SEM of three independent biological experiments, each performed with four technical replicate wells per condition. (C), USA300 was cultured in TSB or TSB supplemented with either 50 µM C16:0 or 20 µM C18:2 as indicated, and cells were harvested for preparation of whole cell lysates after 4h of growth. 20 µg of total protein was subjected to SDS-PAGE and Western blot for detection of FemT as described in B. Brackets represent biological replicates.

### USA300Δ*femT* exhibits impaired growth in presence of exogenous saturated and unsaturated fatty acids

Although production of FemT was not altered in response to exogenous fatty acids (Fig. 1C), its co-expression with *fem*X alludes to a potential role in efflux of lipid substrates, as further implicated through its structural relatedness to SwrC in *B. subtilis*, which promotes efflux of a lipopeptide surfactin. Therefore, to assess the consequence of FemT loss under conditions when stress is imposed on the membrane through exposure to exogenous fatty acids, we constructed USA300Δ*femT*. Loss of FemT expression in USA300Δ*femT* and restoration of expression with pALC*femT* which confers a basal level of expression from the leaky P*_xyl/tet_* promoter-operator was first confirmed by Western blot (Fig. S2). In growth assays, USA300Δ*femT* exhibited impaired growth in TSB + 100 µM linoleic acid C18:2, although the phenotype was less severe compared to loss of *farE* (Fig. 2A), for which expression is induced in response to C18:2, and growth was restored by complementation with pALC*femT* (Fig. 2B). USA300Δ*femT* also exhibited strongly impaired growth in presence of 500µM C16:0 which was ameliorated with pALC*femT* (Fig. 2C). Therefore, although FemT expression was not appreciably affected by exposure to saturated or unsaturated fatty acids, its function is required for optimal growth when *S. aureus* is exposed to these same fatty acids.

**Figure 2.**
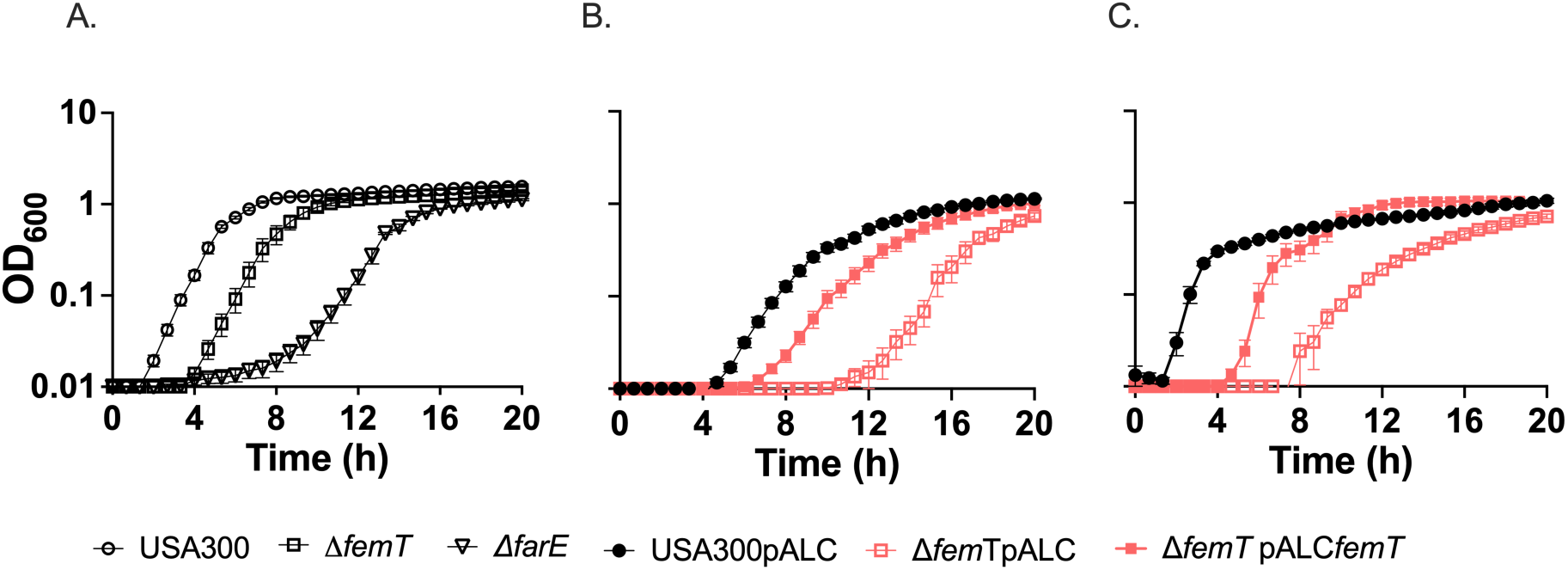
Growth kinetics of USA300, USA300Δ*femT*, and USA300Δ*farE* in presence of unsaturated linoleic acid (A, B) or saturated palmitic acid (C). For panel (A), overnight cultures of USA300, USA300Δ*femT*, and USA300Δ*farE* were diluted to an OD_600_ of 0.01 and inoculated into 96-well plates containing 200 µL TSB supplemented with 100 µM linoleic acid C18:2, and growth was monitored hourly for 20h. (B), Growth kinetics of USA300 + empty pALC vector, USA300Δ*femT* + pALC, and USA300ΔfemT + pALC*femT* in TSB + 100 µM linoleic acid. (C), as in B, except that the culture medium was TSB + 500 µM palmitic acid C16:0. Data represent the mean ± SEM of three independent biological experiments, each performed with six replicate wells per condition.

### USA300Δ*femT* exhibits a strong proteostasis and oxidative stress signature in response to palmitic acid

To better understand how loss of *femT* affects growth in the presence of exogenous fatty acids, we conducted RNAseq analysis of USA300 and USA300Δ*femT* in TSB and TSB + 50 µM palmitic acid, C16:0. We chose 50 µM palmitic acid since it was established to be the highest concentration that was tolerated without substantially affecting growth (data now shown), and saturated palmitic acid also does not induce *farE* expression, allowing more accurate assessment of changes in gene expression attributed to *femT* alone. The complete set of differentially expressed genes is available in Dataset S1, and volcano plots are shown in Fig. 3. In wild type USA300, 50 µM C16:0 promoted increased expression of *fad* genes (Figure 3A), which contribute to metabolism of palmitic acid (Kuiack *et al*., 2023). Surprisingly, there was also significantly increased expression of genes previously known to be induced in response to long chain unsaturated fatty acids, including several genes comprising a Type VII secretion system, as well as cell surface protein SasF implicated in resistance to unsaturated fatty acids (Kenny *et al*., 2009b; Lopez *et al*., 2017a). Oleate hydratase *ohyA* was also significantly enhanced in response to saturated C16:0 even though its enzymatic activity selectively hydroxylates exogenous unsaturated fatty acids (Radka et al., 2021; Subramanian et al., 2019). Significantly down-regulated genes included a small number of cell surface or secreted fibrinogen- and IgG binding proteins (*sbi, vwb, ecb, efb*) and *hla* encoding alpha hemolysin. However, other than a significant increase in expression of *cwrA,* a gene that responds to cell wall damage (Balibar et al., 2010a), there was no indication of a stress response.

**Figure 3.**
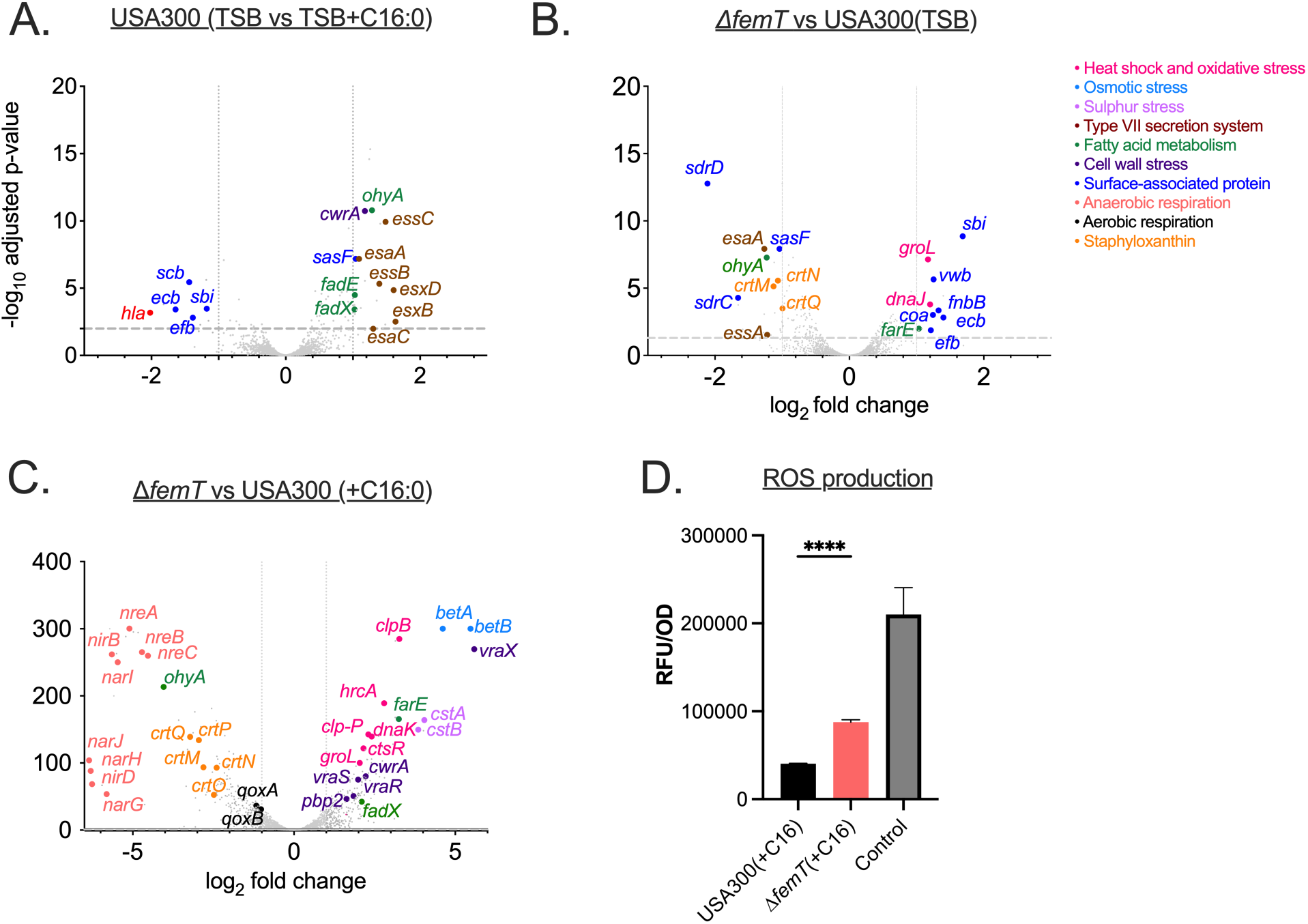
Transcriptomic analysis and physiological validation of oxidative stress in USA300Δ*femT*. Differential gene expression in USA300Δ*femT* compared to USA300 during growth in TSB (A), in USA300 grown in TSB supplemented with 50 µM C16:0 relative to TSB alone (B), and in USA300Δ*femT* compared to USA300 during growth in TSB supplemented with 50 µM C16:0 (C). Overnight cultures were diluted to an OD_600_ of 0.01 in 25 mL TSB or TSB supplemented with 50 µM C16:0 and cultures were grown to OD_600_ ∼0.35-0.4 prior to processing for RNA extraction. The x-axis represents log2-fold change, and the y-axis represents −log10 adjusted p-value. Differential expression thresholds were set at log2-fold change ≥ 1 and adjusted p < 0.01. Selected genes associated with fatty acid metabolism, secretion systems, cell envelope stress, respiration, sulfur stress, and virulence are highlighted, and functional categories are indicated by color coding, as shown in the legend. **(D)** Intracellular reactive oxidative species (ROS) were assessed using the fluorescent probe H_2_DCFDA following growth in TSB supplemented with 50 µM C16:0, or a control culture exposed to 1mM H_2_O_2._ Fluorescence was normalized to cell density. Data represent the mean ± SEM from three biological replicates. Statistical significance was determined using an unpaired two-tailed Student’s *t*-test. ****, *p* < 0.0001.

The reduced expression of *hla* and several genes encoding cell surface or secreted fibrinogen- and IgG binding proteins in response to palmitic acid in USA300 is consistent with prior observations that fatty acids can suppress SaeRS-associated transcription (Ericson *et al*., 2017). Notably, several of these genes (*sbi, vwb, ecb, efb*) exhibited significantly increased expression in USA300Δ*femT* during growth in TSB alone (Figure 3B), potentially indicating altered membrane-responsive signaling through SaeRS in the absence of FemT. Accordingly, the transcriptional signature of USA300Δ*femT* in TSB alone was supportive of a mild stress response, as evident from increased expression of *groEL* and *dnaJ* encoding chaperonins involved in protein folding (Loi *et al*., 2023). Conversely, several genes involved in lipid synthesis, metabolism, or response to host lipids were significantly downregulated including *crt* genes that promote synthesis of carotenoid pigment staphyloxanthin, oleate hydratase (*ohy*A) and the Type VII secretion gene *esa*A, as well as cell surface protein *sas*F (Fig. 3B). Strikingly, the reduced expression of *ohy*A and *crt*QMN noted in TSB alone was much stronger in TSB + 50 µM C16:0 (Fig. 3C), under which condition numerous genes involved in oxygen sensing (*nre*BC) and aerobic and anaerobic respiration (*qox*, *nir* and *nar)* were also strongly down-regulated, while a strong stress response was evident through increased expression of genes involved in heat shock, protein folding and oxidative stress (*msr*A, *hrc*A, *gro*EL, *dna*JK, *clp*BC), sulfur stress (*cst*ABR), cell wall stress and synthesis (*vra*RSTX, *cwr*A, *pbp*2, *sgt*B), and osmotic stress (*bet*AB, *opu*BCD).

USA300Δ*femT* also exhibited a significant increase in expression of *farE*, which was not evident in wild type USA300 exposed to palmitic acid. Therefore, loss of *femT* imparts a mild stress response during growth in TSB alone, accompanied by reduced expression of *ohyA*, *essA* and *sasF* that were previously implicated in the response to or metabolism of unsaturated fatty acids, and reduced expression of *crt* genes involved in synthesis of carotenoid lipid. Conversely, exposure to 50 µM C16:0 imparts a strong stress response characterized by significantly increased expression of genes involved in lipid efflux and metabolism, degradation of misfolded proteins, and oxidative stress and osmo-tolerance, concomitant with strongly reduced expression of genes involved in cellular respiration, as well as *ohy*A and *crt*QMN.

Since an oxidative stress response was manifested in the transcriptional signature of USA300Δ*femT* on exposure to palmitic acid, we tested for accumulation of reactive oxygen species ROS. Consistent with oxidative stress, ROS measurements using H_2_DCFDA revealed significantly greater fluorescence in USA300Δ*femT* than in USA300 following exposure to 50 µM C16:0 (Fig. 3D), providing physiological evidence that loss of FemT results in elevated oxidative stress during fatty acid exposure.

### USA300Δ*femT* exhibits increased membrane rigidity, reduced respiratory activity, and cell division defects on exposure to palmitic acid

Among the strongest effects of C16:0 on gene expression in USA300Δ*femT* were the down-regulation of genes encoding functions required for cellular respiration, accompanied by a dramatic increase in expression of *vraX*, which was previously reported to be induced in response to small molecules that bind and sequester Lipid II (Lu et al., 2023; Zhong et al., 2025). Notably, *vraX* expression also showed an upward trend in USA300Δ*femT* during growth in TSB alone (log2FC-1.6; Dataset S1), although this did not reach statistical significance. For clarification, we constructed a *vraX::lux* reporter, which exhibited strongly increased activity in USA300Δ*femT* relative to USA300 in both TSB and TSB with 50 µM C16:0 (Fig. 4A). Additionally, whereas assays of cellular respiration showed no significant difference between USA300 and USA300Δ*femT* during growth in TSB, USA300Δ*femT* exhibited significantly reduced respiration in TSB + 50 µM C16:0, which was restored with pALC*femT* (Fig. 4B). Reduced respiratory activity was accompanied by loss of membrane potential (Fig 4C), concomitant with increased membrane rigidity (Fig 4D). These data are strongly supportive of a significant perturbation of membrane function in USA300Δ*femT* on exposure to palmitic acid.

**Figure 4.**
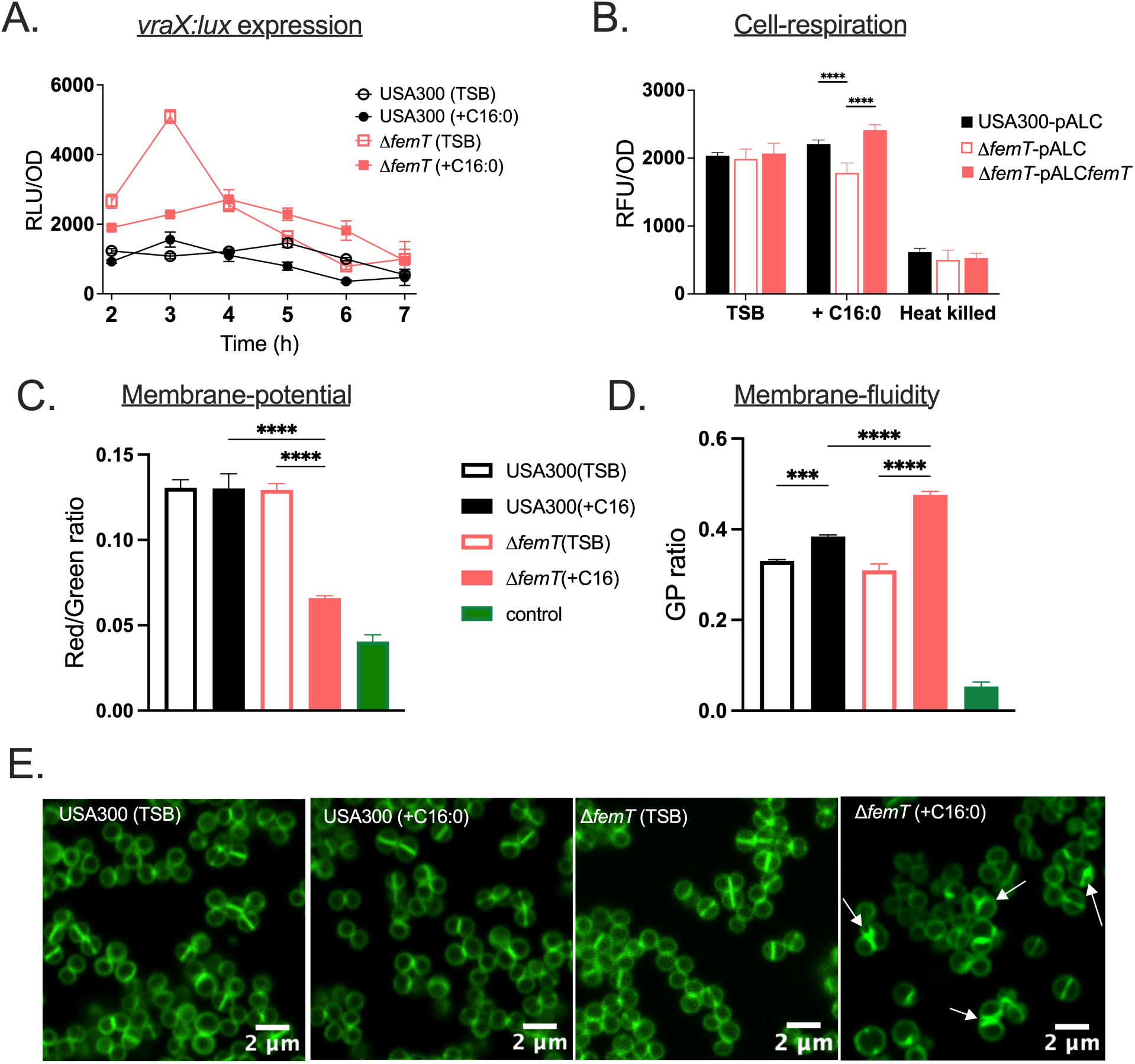
Palmitic acid induces cell wall stress, reduced respiration and membrane potential, altered membrane fluidity, and aberrant cell division in USA300Δ*femT*. (A), expression of *vraX::lux* reporter in USA300 and USA300Δ*femT* during growth in TSB or TSB + 50 µM C16:0; (B), cellular respiration measured by CTC reduction assay in USA300 pALC, USA300Δ*femT* pALC, and USA300Δ*femT* pALC*femT*; (C,D) membrane potential measured using DiOC_2_(3) fluorescence and membrane fluidity assessed using Lauran dye in USA300 and USA300Δ*femT* grown in TSB or TSB supplemented with 50 µM C16:0; (E) fluorescence microscopy of cell wall and septal morphology using BODIPY-vancomycin staining. For *vraX*::*lux* reporter assays (A), cultures of USA300 and USA300Δ*femT* harboring *vraX::lux* were diluted to an OD_600_ of 0.01 in 25 mL TSB and incubated at 37 °C with shaking. At hourly intervals, culture aliquots were removed for OD_600_ determination, and separate 4 x 200 µl aliquots were transferred to 96 well microtiter plates for luminescent detection. Data are expressed as relative light units [RLU]/OD_600_. Cellular respiration (B) was measured using the CTC reduction assay in USA300 pALC, USA300Δ*femT* pALC, and USA300Δ*femT* pALC*femT* grown in TSB or TSB + 50 µM C16:0. Heat-killed/control samples were included as a negative control. Fluorescence was normalized to cell density (RFU/OD_600_). Membrane potential (C) measured using DiOC_2_(3) fluorescence in USA300 and USA300Δ*femT* grown in TSB, TSB supplemented with 50 µM C16:0. Where indicated, cells were treated with gramicidin D (Sigma-Aldrich) at a final concentration of 1 μg/mL for 5 min prior to analysis and served as a depolarized control. Membrane potential is expressed as the red/green fluorescence ratio. Membrane fluidity (D) assessed using Laurdan generalized polarization (GP) in USA300 and USA300Δ*femT* grown in TSB or TSB supplemented with 50 µM C16:0 to mid-exponential phase. Cultures treated with 100 µM linoleic acid for 15 min at 37°C prior to Laurdan staining served as a membrane fluidization control. Higher GP values indicate increased membrane order (rigidity), whereas lower values indicate increased membrane fluidity. Fluorescence microscopy (E) of USA300 and USA300Δ*femT* grown in TSB or TSB supplemented with 50 µM C16:0 to an OD_600_ of ∼0.3–0.4. Cells were washed, fixed, and stained with BODIPY-vancomycin, and imaged using confocal microscopy. Arrows indicate aberrant septa, including asymmetric and Y-shaped division planes. Scale bars, 2 µm. Each data point represents the mean ± SEM of three independent biological replicates, each performed using triplicate flask cultures. Statistical significance was determined using two-way ANOVA with Tukey’s multiple-comparisons test. Significance is indicated as follows: *p* < 0.05(*); *p* < 0.01(**); *p* < 0.001(***); *p* < 0.0001(****). Control samples were excluded from statistical comparisons.

To assess whether altered membrane properties were reflected in cell morphology, we examined cell division and septation using BODIPY-vancomycin labeling. USA300 and USA300Δ*femT* grown in TSB alone, and USA300 grown in TSB + 50 µM C16:0 all exhibited similar morphology with clearly defined division septa, whereas USA300Δ*femT* in TSB + 50 µM C16:0 exhibited heterogeneity in cell size and aberrant division septa, including Y-shaped and asymmetric septa (Fig. 4E, S3). Additionally, quantitative analysis revealed increased variability in cell volume among USA300Δ*femT* cells grown in TSB supplemented with 50 µM C16:0 compared to USA300 (Figure S4). Collectively, these findings demonstrate that loss of FemT under palmitic acid conditions is associated with activation of a cell envelope stress response, impaired respiratory and membrane function, and defects in cell division and septum formation.

### USA300Δ*femT* exhibits a significantly altered metabolome in both TSB and TSB + 50 µM palmitic acid

For further insight into the stress response inherent in the RNAseq data and altered respiratory activity, we conducted metabolomics analyses of cultures grown in TSB and TSB + 50 µM C16:0. A complete list of all detected metabolites, including raw and normalized abundance values and the input data used for statistical analyses, is provided in Dataset S2, while a heat map of the top 30 differentially affected metabolites is shown in Fig. 5. Although one replicate for USA300 in each condition behaved anomalously, the remaining replicates in these two groups were more similar to one another than to USA300Δ*femT* grown in TSB, while the metabolite pool of USA300Δ*femT* grown in TSB + 50 µM palmitic acid was most divergent. For USA300 grown with palmitic acid compared to TSB alone, the most striking differences were elevated urocanate, glutamate and cystathionine. Urocanate is indicative of histidine catabolism with release of ammonia through the *hut* pathway, with glutamate as an end-product (Bender, 2012). Concurrent elevation of urocanate, glutamate and cystathionine in response to palmitic acid may infer a requirement for increased pH and redox buffering capacity (Lo et al., 2009; Soutourina et al., 2010; Beetham et al., 2024; Liechti and Goldberg, 2026), reflecting a routine physiologic response to exogenous fatty acids.

**Figure 5.**
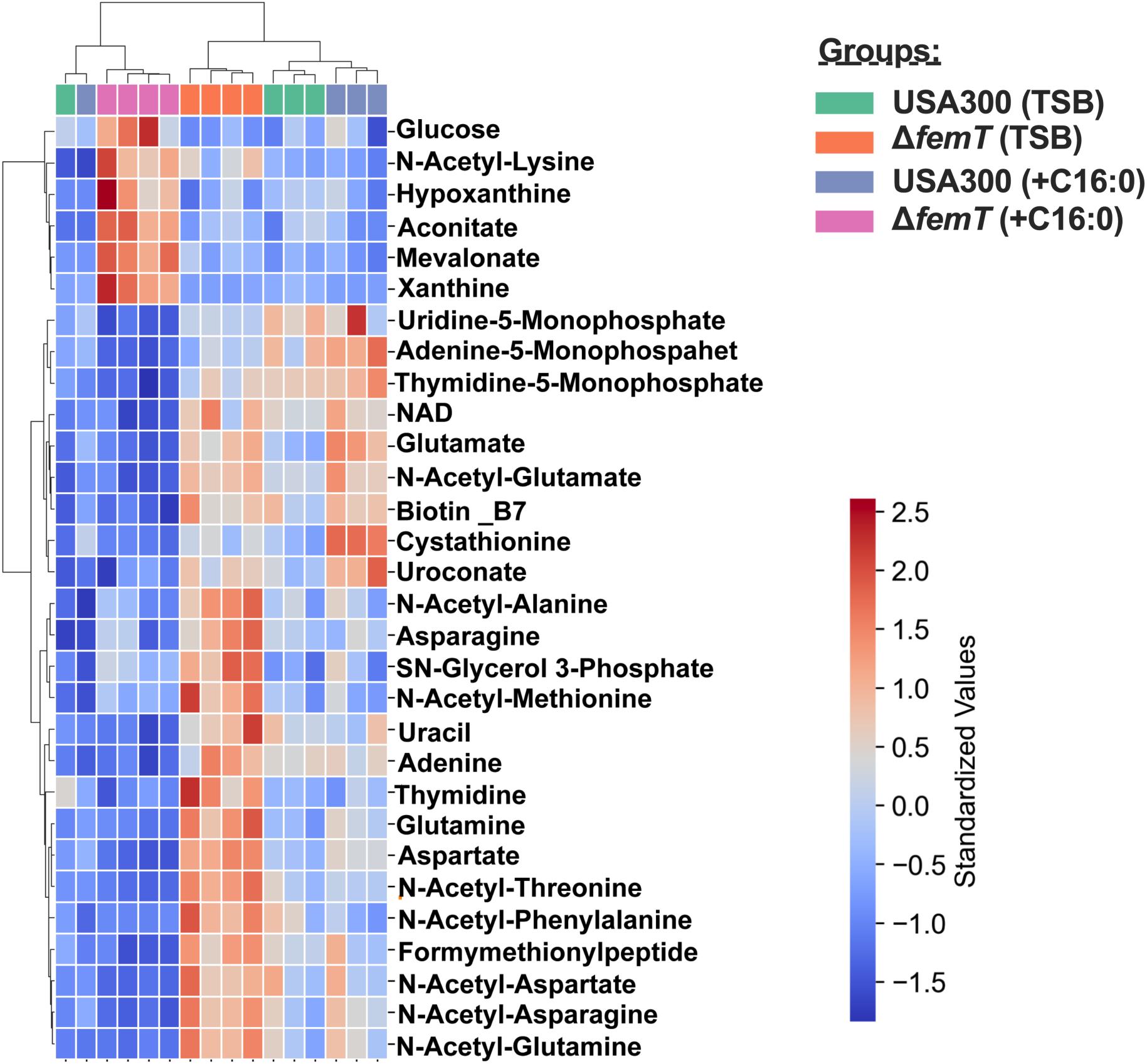
Heat map of top 30 differentially affected cellular metabolites in USA300 and USA300Δ*femT* during growth in TSB or TSB supplemented with palmitic acid (C16:0). Overnight cultures were diluted to an OD_600_ of 0.01 in TSB with or without 50 µM C16:0 and grown at 37 °C with shaking to OD_600_∼0.35–0.4 prior to metabolite extraction. Heat maps were generated from auto-scaled metabolite intensities. Significant features were identified by one-way ANOVA with Benjamini–Hochberg false discovery rate (FDR) correction and ranked by adjusted p-value. Data were scaled row-wise, and hierarchical clustering was performed using Euclidean distance and Ward linkage. Data represent four biological replicates per condition. Red and blue indicate higher and lower abundance, respectively.

Comparison of cultures grown in TSB alone revealed elevated glutamine, aspartate and asparagine in USA300Δ*femT*, which may reflect reduced incorporation of nitrogen into biomass or macromolecules. Evidence of stress is inherent in elevated levels of several acetylated amino acids, which may reflect degradation of oxidized proteins (Rizo and Encarnación-Guevara, 2024), while elevated sn-glycerol 3-phosphate is suggestive of membrane remodeling or altered energetics (Liu *et al*., 2022). Remarkably, most of the metabolites that were elevated during growth of USA300Δ*femT* in TSB were strongly diminished in TSB + 50 µM C16:0, under which condition there was elevated N-acetyl-lysine, which may indicate increased protein acetylation and/or proteolysis, and can also reflect carbon excess relative to growth capacity (Nakayasu *et al*., 2017), which is further supported by elevated glucose. Altered carbon flux is further evident in accumulation of aconitate, reflecting reduced carbon flow into the TCA cycle, in addition to oxidative or nitrosative stress (Somerville *et al*., 2002) while accumulation of mevalonate is indicative of oxidative and energy stress as well as active membrane or cell wall remodeling (Balibar et al., 2008). Consistent with these changes, NAD⁺ levels were markedly reduced in Δ*femT* under palmitic acid conditions (Figure 5), further supporting disruption of cellular redox and energy homeostasis. Lastly, strongly elevated hypoxanthine and xanthine is a high confidence signature of purine degradation with impaired nucleotide salvage (Lithgow et al., 2004). These data are supportive of a mild oxidative stress in USA300Δ*femT* during growth in TSB alone, which is strongly exacerbated on exposure to 50 µM C16:0, leading to profound metabolic remodeling.

### Disruption of *femT* affects cellular lipid composition

Our data thus far are indicative of oxidative stress, metabolic reprogramming, altered energetics and membrane remodeling in USA300Δ*femT* on exposure to palmitic acid. We therefore conducted lipidomic analyses for further insight. For this purpose, we chose oleic acid (C18:1) which is normally well tolerated by *S. aureus* and can be easily monitored since *S. aureus* does not synthesize unsaturated fatty acids.

A comprehensive list of annotated lipid species, including their identification parameters and normalized abundance values across all conditions, is provided in Dataset S3. Analysis of cellular free fatty acids revealed that C18:1 was not evident in TSB alone but was abundant in cultures supplemented with C18:1 (Fig. 6A). Lesser levels of C20:1 in both USA300 and USA300Δ*femT* indicate that exogenous C18:1 was being actively metabolized and extended to C20:1 via the FASII cycle (Fig 6B). Despite this, 10-hydroxystearic acid which is a metabolite produced by the action of oleate hydratase OhyA on C18:1 was elevated only in USA300 (Fig 6C). This is consistent with transcriptomics data where *ohyA* expression was significantly reduced in USA300Δ*femT* in TSB alone and strongly diminished in TSB + 50 µM C16:0.

**Figure 6.**
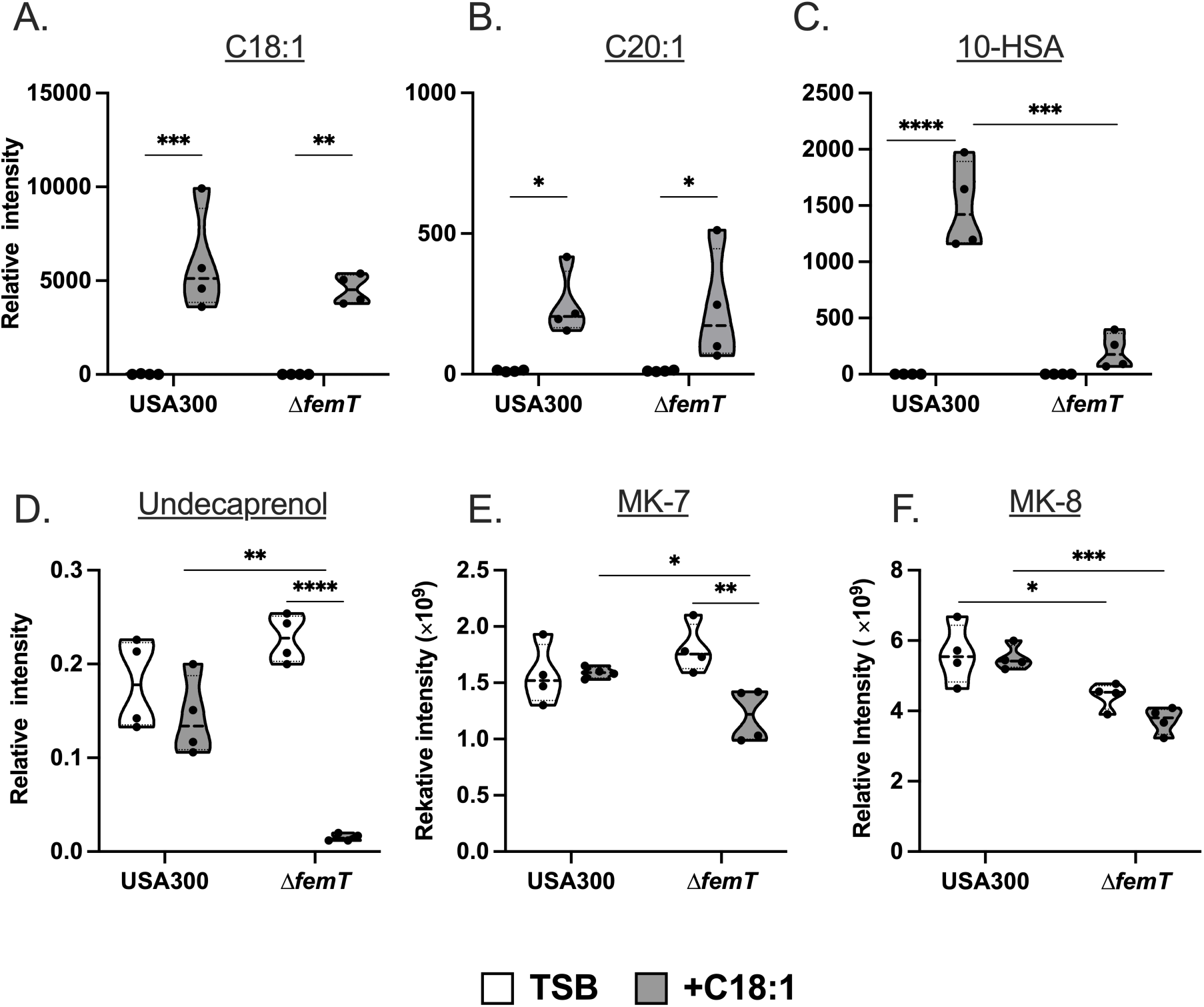
Lipidomics comparison of selected free fatty acids, undecaprenol and menaquinones in USA300 and USA300Δ*femT*. Relative abundance of fatty acid–derived species in USA300 and USA300Δ*femT* grown in the presence of exogenous oleic acid (C18:1), including C18:1 (A), C20:1 (B), and 10-hydroxystearic acid (10-HSA) (C), as well as undecaprenol (D) and menaquinone species MK-7 (E) and MK-8 (F). Prior to processing for lipid extraction, mid-exponential phase cultures (OD_600_ ∼0.4) grown in TSB + 0.1% DMSO were supplemented with either 500 µM C18:1 or equivalent volume of solvent sham (0.1% DMSO in TSB), followed by continued growth to OD_600_ of ∼1.0. Data are shown as individual data points with violin plots indicating distribution and represent four biological replicates (independent flasks) per condition. Statistical significance was determined using two-way ANOVA with Tukey’s ‘multiple comparisons. *p* < 0.05(*); *p* < 0.01(**); *p* < 0.001(***); *p* < 0.0001(****).

The data also revealed strongly diminished production of C55 isoprenoid (undecaprenol) lipid carrier in USA300Δ*femT* grown with exogenous C18:1 compared to USA300 (Fig. 6D). A less severe but still significant reduction was also noted for the isoprenoid menaquinones MK-7 and MK-8 (Fig’s 6E, F), which participate in electron transport and oxidative phosphorylation (Götz and Mayer, 2013; Desai *et al*., 2016). The major phosphatidylglycerol (PG) species during growth in TSB were PG 33:0, PG 34:0 and PG 35:0 with no significant differences between USA300 and USA300Δ*femT*, and each of these were reduced in abundance when cells were grown in TSB + C18:1, but with no significant differences between strains (Fig 7A). The reduced abundance of these major PG species in response to C18:1 was accompanied by appearance of new PG species that incorporated C18:1 or its FASII extension products. Of these, USA300Δ*femT* exhibited significantly increased abundance of PG 33:1 and PG 35:1 compared to USA300 (Fig 7B), in addition to less abundant PG’s that were unique to USA300Δ*femT* and distinguished by having two unsaturated fatty acids, including PG 18:1_18:1, PG 18:1_20:1, and PG 18:1_22:1 (Fig 7C). To investigate how C18:1 affects membrane order, we conducted a laurdan dye general polarization assay (laurdan GP) where a high ratio reflects tightly packed lipids with low hydration and a low ratio indicates a more loosely packed less ordered membrane. USA300Δ*femT* grown with 50 µM C18:1 exhibited a significantly less ordered membrane relative to wild type USA300 (Fig S5). Together, these findings indicate that loss of FemT leads to enhanced incorporation of oleic acid into membrane phospholipids, leading to a less ordered membrane and widespread disruption of membrane lipid homeostasis.

**Figure 7.**
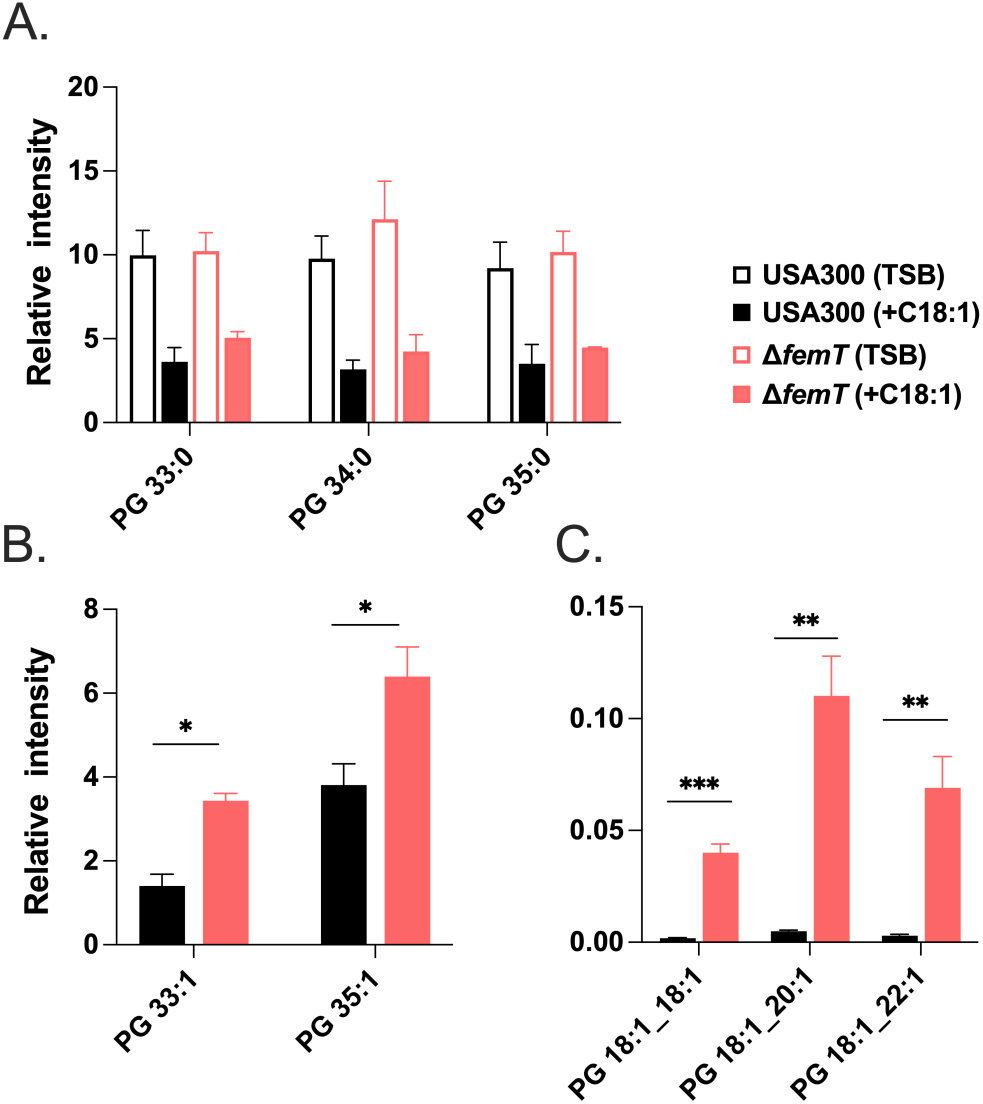
Differential incorporation of oleic acid C18:1 and its FASII extension products (C20:1 and C22:1) into phosphatidylglycerol PG of USA300 and USA300Δ*femT*. Relative intensity of individual phosphatidylglylcerol (PG) species is derived from lipidomics data. (A), major PG species derived from endogenously synthesized fatty acids in USA300 and USA300Δ*femT* grown in TSB, or TSB with bolus of 500 µM C18:1. The mass values of individual PG species correspond to fatty acid profiles PG33:0, C15:0 + C18:0; PG 34:0, C15:0 + C19:0; and PG 35:0, C15:0 + C20:0. (B), relative abundance of major PG species containing exogenous C18:1 or its FASII extension product C20:1 from cells supplemented with C18:1; PG 33:1, C15 + C18:1; PG 35:1, C15 + C20:1. (C), relative abundance of minor PG species containing two unsaturated fatty acids that were unique to USA300Δ*femT* cells supplemented with exogenous C18:1; PG 18:1_18:1, PG 18:1_20:1 and PG 18:1_22:1 as indicated, containing oleic acid C18:1 or its FASII extension products C20:1 and C22:1. Data are represented as mean relative intensity ± SEM from four biological replicates. Statistical significance was determined using two-way ANOVA with Tukey’s multiple-comparisons test. Significance is indicated as follows: *p* < 0.01 (**), *p* < 0.001 (***), and *p* < 0.0001 (****).

### USA300Δ*femT* exhibits attenuated virulence in a murine skin abscess infection model

Given that USA300Δ*femT* exhibited signatures of oxidative and osmotic stress in response to palmitic acid, a fatty acid abundant in host environments such as the skin, we next sought to assess the *in vivo* relevance of this phenotype using a subcutaneous skin infection model. To test this, C57BL/6 mice were subjected to subcutaneous flank infection with either WT USA300 or USA300Δ*femT* and monitored for weight loss, lesion development, and bacterial burden at the infection site. Mice infected with USA300Δ*femT* developed moderately smaller lesions compared to those infected with WT, and this reduction was statistically significant (Figure 8A). Additionally, USA300Δ*femT*-infected mice exhibited significantly lower percentage weight loss as compared to USA300 infected mice (Figure 8B). However, bacterial counts at lesion sites were similar between WT and USA300Δ*femT* (Figure 8C). Although the impact on virulence was modest, these data are supportive of FemT contributing to virulence in *S. aureus*.

**Figure 8.**
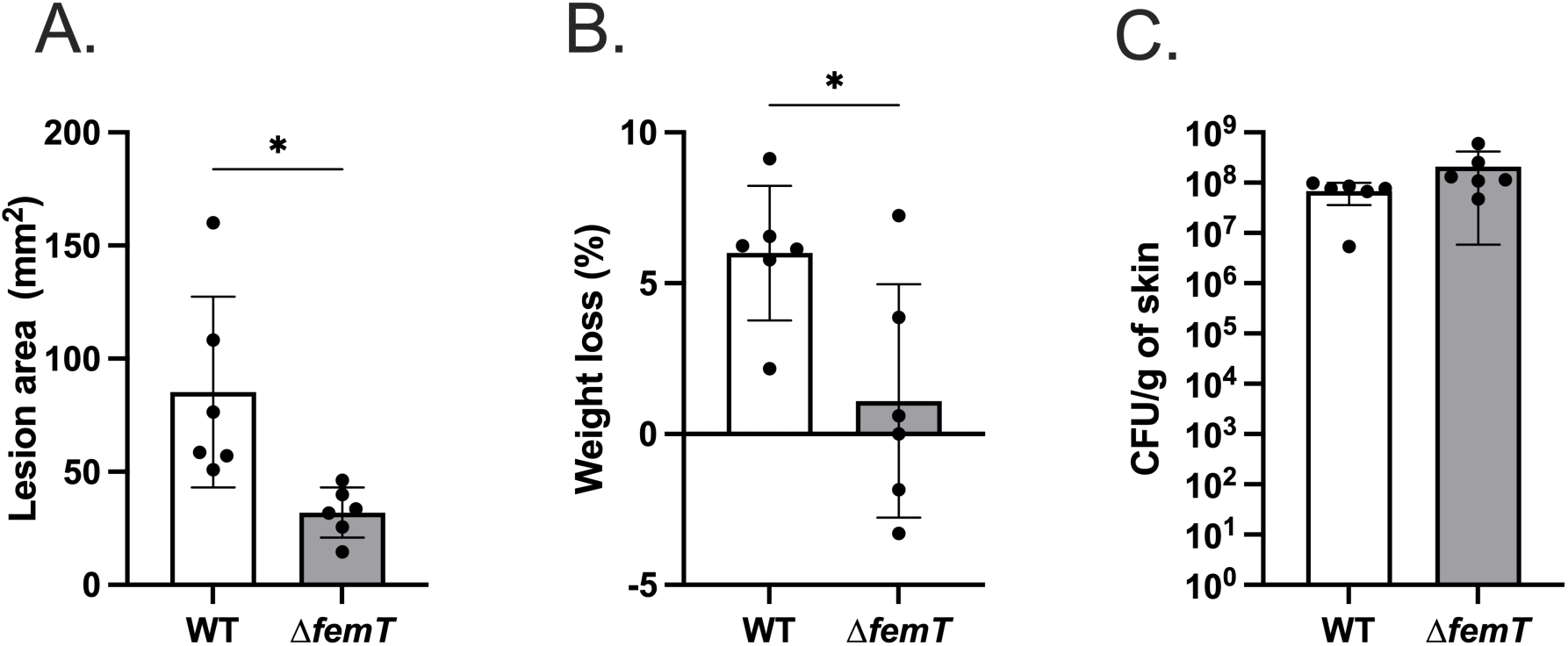
Loss of FemT attenuates lesion severity during *Staphylococcus aureus* skin infection. Lesion area (A), percentage of body weight loss (B), and bacterial burden in infected skin tissue (C) in mice infected with USA300 or USA300Δ*femT*. Eight- to twelve-week-old male and female C57BL/6 mice were infected subcutaneously in the shaved lower flank with ∼5 × 10⁷ CFU of *S. aureus* USA300 (WT) or USA300Δ*femT* suspended in Hanks’ balanced salt solution. Lesion area was measured at day 3 post-infection, and values represent the average lesion size per mouse (A). Percentage body weight loss was measured at day 3 post-infection (B). Bacterial recovery from infected skin tissue at day 3 post-infection is expressed as CFU per gram of tissue following lesion excision, homogenization, and serial dilution plating (C). Each point represents an individual mouse, and bars indicate mean ± SE. Statistical significance was determined using an unpaired two-tailed Student’s *t* test. Significance is indicated as follows: *p* < 0.05(*); *p* < 0.01(**).

## DISCUSSION

This study provides the first detailed insight into the function of the RND efflux pump FemT in *S. aureus*, which is co-expressed with *femX*. Based on FemX having an essential role in synthesizing the Lipid II precursor of peptidoglycan, an important accessory function for FemT in peptidoglycan synthesis was envisioned. Nevertheless, inactivation of *femT* was well tolerated in the absence of imposed stress, and our initial data were not supportive of a role for FemT in lipid homeostasis, since its constitutive expression was not altered in response to either saturated or unsaturated fatty acids, and despite its expression being dependent on its co-association with *femX*, inactivation of *femT* function did not cause any growth impairment in the absence of an imposed stress. However, our implementation of a suite of transcriptomic, metabolomic and lipidomic tools allowed us to identify key traits in USA300Δ*femT* during unimpaired growth in TSB, which became profound in the presence of palmitic acid. This included expression of FarE in response to palmitic acid in USA300Δ*femT* but not in wild type USA300, and FarE has previously been implicated in efflux of phosphatidylglycerol and host lipid sphingosine (Chen *et al*., 2022). Cumulatively, our data are strongly supportive of a role for FemT in maintaining cellular lipid homeostasis, leading to compensatory expression of FarE in response to loss of FemT function.

Although USA300Δ*femT* exhibited no growth defect in TSB alone, transcriptomic data revealed significantly reduced expression of *ohyA* and *crtMN* encoding enzymes that modify lipid substrates; OhyA functioning to detoxify oleic acid and presumably other unsaturated fatty acids through hydroxylation of double bonds, while CrtMN catalyze initial steps in conversion of farnesyl pyrophosphate into the carotenoid pigment Staphyloxanthin (Pelz *et al*., 2005). This was accompanied by significantly increased expression of GroEL and DnaJ that aid in re-folding of misfolded or damaged proteins. Consistent with this proteostasis response, metabolite analysis revealed accumulation of acetylated amino acids, likely reflecting degradation of carbonylated proteins (Chatterjee *et al*., 2011). Together, these data suggest that even in the absence of exogenous fatty acid, loss of FemT results in altered lipid homeostasis and an elevated basal stress state eliciting proteostasis and metabolic adaptations.

Strikingly, expression of *ohyA* and *crt* were significantly reduced during growth of USA300Δ*femT* in TSB alone and profoundly reduced in response to C16:0. In sharp contrast, wild type USA300 exhibited strongly enhanced expression of *ohyA* in response to C16:0 which was unexpected since this enzyme selectively hydroxylate unsaturated fatty acids (Radka, Batte, Frank, Young, et al., 2021). Several Type VII secretion system genes that were previously noted for increased expression in response to unsaturated linoleic acid C18:2 (Lopez et al., 2017) also showed strongly increased expression in response to saturated C16:0. Therefore, increased expression of *ohyA* and Type VII secretion system genes seems to be a general response to exogenous fatty acids irrespective of their saturation status. Although OhyA is known for hydroxylating unsaturated fatty acids (Radka, Batte, Frank, Rosch, et al., 2021; Subramanian et al., 2019) it may act on other lipid classes including isoprenoids, terpenoids and carotenoids such as Staphyloxanthin, all of which have double bonds that would be susceptible to oxidation. Since our data also revealed elevated ROS in USA300Δ*femT* on exposure to palmitic acid, the strongly reduced expression of *ohyA* and *crt* under this condition likely reflects a compensatory mechanism to limit the accumulation of oxidized lipids.

Additional evidence of this protective response is inherent in the elevated mevalonate in USA300Δ*femT* in response to 50 µM C16:0 (Figure 6). The mevalonate pathway generates isopentenyl pyrophosphate, which is a precursor to synthesis of terpenoids, carotenoids and quinols, including the undecaprenol lipid carrier C55P, Staphyloxanthin pigment, and menaquinones required for respiration (Misic *et al*., 2016), all of which are prone to oxidative damage. The noted accumulation of mevalonate is suggestive of a blockade in synthesis of the various terpenoid, carotenoid and quinone end products. This is supported by our lipidomics analysis of USA300Δ*femT* exposed to C18:1, which revealed a strong reduction in undecaprenol required for cell envelope biosynthetic reactions, and a significant reduction in the cellular pool of MK-7 and MK-8, which are the major menaquinones used for oxidative phosphorylation in *S. aureus* (Collins and Jones, 1981; Götz and Mayer, 2013).

Another striking phenotype of USA300Δ*femT* grown in TSB with 50 µM C16:0 was accumulation of N-acetyl lysine. On a general level, accumulation of acetylated amino acids likely reflect degradation of post-translationally modified proteins, as supported by the strongly enhanced expression of the *ctsR* and *hrcA* regulators of proteostasis, and the ClpP protease which degrades damaged proteins (Lund, 2013). However, specific and reversible N-acetylation of lysine has also been noted a means of modulating the activity of microbial acetyl-CoA synthase, whereby lysine acetylation inactivates the enzyme, and activity is restored through a sirtuitin-like deacetylase (Starai and Escalante-Semerena, 2004; Crosby et al., 2012; Burckhardt, Buckner and Escalante-Semerena, 2019). Acetylome profiling in the diatom *Phaeodactylum tricornutum* also revealed extensive lysine acetylation of proteins involved in fatty acid metabolism (Chen *et al*., 2018), including multiple lysine acetylation of a long chain acyl-CoA synthase which has a fundamental role in providing acyl-CoA for fatty acid and phospholipid synthesis, and fatty acid degradation through β-oxidation. However, in this example lysine acetylation stimulated activity, rather than inhibition that was noted with microbial acetyl-CoA synthases. While we do not have direct proof of proteins with acetylated lysine in USA300Δ*femT* cells, the accumulation of N-acetyl lysine is a strong indicator of this process, which is known to inactivate acetyl-CoA synthase activity in bacteria and stimulate long chain acyl-CoA synthase activity in diatoms.

A limitation of our work is that the data are correlative to loss of FemT function in cells stressed with exogenous fatty acids, but this has not yet revealed a physiologic efflux substrate. This is not unusual for studies on RND efflux pump function, where years of study combined with appropriate analytic tools have come to reveal a general role in metabolite efflux and modulation of stress responses (Yamasaki et al., 2023). Accordingly, the requirement for FemT during growth under imposed stress mirrors the prototypic RND efflux pump AcrB in *E. coli*, first described for its role in resistance to acridine dye and being induced in response to stress conditions (D. Ma et al., 1995; Nakamura et al., 1978). More recent studies on AcrB in *Salmonella* revealed that its inactivation led to altered membrane potential, accompanied by increased and prolonged expression of genes responsible for anaerobic energy metabolism (Whittle *et al*., 2024), which was proposed to be linked to sensing of an altered redox state of the cellular quinol pool through the ArcAB TCS of oxidative stress. Remarkably, the transcriptome of USA300Δ*femT* grown with 50 µM C16:0 appears to phenocopy a deletion mutation in the *srrAB* TCS of *S. aureus* which in response to nitrosative stress promotes expression of *qoxAB* required for cytochrome synthesis, as well as *pflAB*, *adhE*, *nrdDG* which contribute to anaerobic metabolism, *scdA* iron-sulfur cluster repair protein and *hmpA* which contributes to nitric oxide detoxification (Kinkel *et al*., 2013), all of which exhibited significantly reduced expression in USA300Δ*femT* exposed to 50 µM C16:0. In view of it being thought that SrrAB senses impaired electron flow through electron transport pathways, it is additionally noteworthy that USA300Δ*femT* exposed to C16:0 also exhibited a profound reduction in expression of the *nreABC* regulator of nitrate respiration, as well as the co-associated *nir* and *nar* genes encoding nitrite and nitrate reductases Therefore, numerous genes that are regulated through the *srrAB* and *nreABC* TCS of aerobic and anaerobic respiration exhibit diminished expression in USA300Δ*femT* cells exposed to subinhibitory C16:0.

From these considerations, it is evident that just as with Gram negative Salmonella where inactivation of AcrB led to altered membrane potential and aberrant expression of genes responsible for energy metabolism, a similar response is noted with loss of FemT function in *S. aureus*. It is therefore salient to reconsider that FemT in *S. aureus* together with SwrC in *B. subtilis* have predicted Alphafold structures that closely resemble Gram-negative RND efflux pumps, for which a key mechanistic tenet is their interaction with accessory proteins to promote a contiguous channel that bridges the inner and outer membrane. Indeed, consistent with its co-expression with *femX*, FemT was found to interact with peptidoglycan biosynthetic enzymes including FemB and PBP2 (Quiblier et al., 2013), while SwrC was found to interact with an accessory lipoprotein MeeY required for surfactin export (He *et al*., 2025). Therefore, although the cell envelope architecture of Gram-positive bacteria does not necessitate that their RND efflux pumps interact with accessory proteins to facilitate export, the structural relatedness of FemT and SwrC to Gram negative RND efflux pumps may reflect a common requirement for interaction with accessory proteins.

As a unifying hypothesis for FemT function, we propose that isoprenoids, terpenoids, carotenoids and menaquinones, all of which have multiple unsaturated carbon bonds, are prone to oxidative damage and must be removed to maintain unimpaired membrane function. As such, co-expression of *femT* with *femX* which is needed to complete the synthesis of the Lipid II precursor of peptidoglycan would reflect a housekeeping function in efflux of oxidized lipid substrates that arise as a consequence of routine metabolic activity. Notably, menaquinones cycle between reduced and oxidized forms during electron transport, while C55P exists in equilibrium with C55PP, which upon delivery of biosynthetic precursors across the cytoplasmic membrane must be dephosphorylated to facilitate recycling to the cytoplasm for a new round of synthesis. It is feasible that through multiple recycling events the C55 backbone becomes oxidized, and menaquinones are also susceptible to oxidative damage through their cyclic reduction and oxidation in electron transport, such that a physiologic function of FemT would be to efflux oxidized or otherwise damaged lipids and lipid metabolites, including hydroxylated fatty acids, C55 and menaquinones. This is consistent with previously proposed functions for RND efflux pumps in Gram-negative bacteria, which includes efflux of fatty acids that are replaced due to membrane damage or phospholipid turnover (Fraud et al., 2008; Adebusuyi and Foght, 2011).

## MATERIAL AND METHODS

### FarE and FemT Structural Comparisons

Protein structures for FemT and FarE from *Staphylococcus aureus* USA300 were obtained from the AlphaFold Protein Structure Database (Varadi *et al*., 2022). The AlphaFold model identifier for FemT (AF-A0A0H2XER4-F1) and additional models used in this study are listed in Table S1. To identify structurally related proteins, the FemT structure was submitted to the DALI server (Holm, 2020), which detects structural homologs based on three-dimensional similarity. Hits were ranked according to Z-score, RMSD, and alignment length. Based on prior literature indicating sequence similarity between FemT and the RND transporter SwrC (SrfP/YerP) from *Bacillus subtilis* (Tsuge, Ohata and Shoda, 2001), pairwise structural comparisons were performed between FemT and SwrC using the DALI server. Structural alignments and superimpositions were visualized using PyMOL (Schrödinger, LLC).

### Bacterial strains and growth conditions

*S. aureus* and *Escherichia coli* strains and plasmids that were used or constructed for this study are listed in Table S2. Strains were stored at −80°C in 20% glycerol and streaked on either Luria–Bertani (LB) agar (*E. coli*) or TSB agar (TSA) plates (*S. aureus*) containing 15 g/L agar when needed for experimental purposes. Tryptic soy broth (TSB) containing 2.5 g/L glucose or TSB without glucose were supplied by Bacto, while LB was purchased from Sigma-Aldrich. For plasmid maintenance, TSA or TSB was supplemented with chloramphenicol at 5 μg/mL, while LB was supplemented with 100 μg/mL ampicillin. Where indicated, *S. aureus* cultures were supplemented with saturated or unsaturated fatty acids. Saturated palmitic acid (C16:0) was obtained from Cayman Chemicals, while unsaturated fatty acids linoleic acid (C18:2) or oleic acid (C18:1) were obtained from Sigma. To supplement media with unsaturated oleic or linoleic acid, a 5 mM stock was first prepared in TSB containing 0.1% dimethyl sulphoxide (DMSO) and then diluted into TSB or warm TSA plus 0.1% DMSO to achieve the desired concentration of fatty acids. To supplement cultures with saturated palmitic acid, a 100 mM stock was first prepared in 70% ethanol and then diluted in TSB 0.1% DMSO to achieve the final desired concentration. Prior to use for phenotypic assays, inoculum cultures were prepared by selecting single colonies of *S. aureus* from agar plates, which were inoculated into 3 mL of TSB in a 13 mL polypropylene tube containing antibiotics as required, and grown at 37°C for 16 h. Unless otherwise indicated, all cultures were grown at 37°C with orbital shaking set at 200 rpm.

### Strain and Plasmid construction

Genetic manipulation of *S. aureus* was conducted in accordance with established guidelines in previous work (Novick, 1991; Arsic *et al*., 2012; Alnaseri *et al*., 2019; Kuiack *et al*., 2023). Restriction enzymes and T4 DNA ligase were purchased from New England BioLabs, Taq polymerase from GenScript, kits for PCR cleanup and plasmid preparation from Genaid, and oligonucleotide primers from Integrated DNA Technologies. All recombinant plasmids were initially constructed in *E. coli* DH5α and their integrity was confirmed by sequencing of the cloned DNA fragment. All shuttle vectors were then transformed by electroporation into *S. aureus* USA300 or isogenic derivatives. Primers used for PCR amplification are listed in Table S3.

Deletion mutants of *femT* and *farE* in the USA300 background were generated using the allelic exchange vector pKOR1 as previously described (Bae and Schneewind, 2006). For construction of the Δ*femT* mutant, ∼1 kb fragment upstream and downstream of *femT* were amplified using primer pairs *femT*UP_*attB1* /*femT*UP_*SacII* and *femT*DW_*SacII* / *femT*DW_*attB2* respectively. The upstream and downstream segments were digested with *SacII* and ligated together for incorporation into pKOR1 using BP Clonase II (Invitrogen). The resulting pKORΔ*femT* was transformed into USA300 and subjected to two-step temperature shift and antisense counterselection, generating USA300Δ*femT.* The *ΔfarE* deletion construct was generated using the same strategy, employing primers *farE*UP_*attB1* and *farE*UP_*SacII* for amplification of a *farE* upstream segment, and *farE*DW_*SacII* and *farE*DW_*attB2* for a downstream flanking segment. Successful construction of the Δ*femT* and Δ*farE* mutants was confirmed by PCR and DNA sequencing.

Plasmid pGYlux (Mesak et al., 2009) was used to create reporter gene constructs for *femT, femX* and *vraX*. Primers *femT::lux_F* and *femT::lux_R* were used to amplify the intergenic region between *femX* and *femT* using USA300 LAC DNA as a PCR template. The product was digested with *Xma*I and *Sal*1-HF, then ligated to *Xma*I/*Sal*I digested pGYlux with T4 DNA ligase to construct pGY*femXT::lux*. Similarly, using primers *femX::lux*_F/*femX::lux*_R and *vraX::lux*_F/*vraX::lux*_R promoter regions of *femX* and *vraX* were amplified respectively and cloned into pGYlux. The complementation plasmid pALC2073 (Corrigan & Foster, 2009) which provides a basal level of gene expression from the P*_xyl/tet_* promoter and inducible expression with anhydrotetracycline was used for ectopic expression of *femT.* The *femT* coding sequence and ribosome binding site was amplified using primer pair pALC*femT*_F/pALC*femT*_R and digested with *Sac*I, ligated with T4 DNA ligase and cloned into the pALC2073, and plasmids with insert were screened by plasmid sequencing to confirm the correct orientation and integrity of *femT* in the pALC*femT* construct. Constructs for expressing the soluble extracytoplasmic transporter domains of FemT were generated using splice overlap extension (SOE) PCR as described previously (Horton *et al*., 1993). FemT has 12 transmembrane helices with two extracytoplasmic soluble domains, SD1 located between helices 1–2, and SD2 between helices 7 and 8. SD1 and SD2 were amplified separately from *S. aureus* genomic DNA using primer pairs of *femT*-

SD1_F/*femT*-SD1_R and *femT*-SD2_F/*femT*-SD2_R, respectively (Table S3). The amplified fragments SD1 and SD2 fragments were gel purified and mixed in equimolar ratios, then fused by overlap extension PCR using Q5 High-Fidelity DNA polymerase (NEB) with 200μM dNTPs, through 15 cycles of denaturation at 95°C for 1 min, annealing at 50°C, and extension at 72°C for 2 min. The outer primer pair *femT*-SD1_F/*femT*-SD2_R was then added to the PCR mixture, followed by another 20 cycles of PCR (denaturation at 95°C for 1 min, annealing at 50°C and extension at 72°C for 2 min). The fused SD1 and SD2 were then gel purified, digested with *NheI* and *SalI* and cloned into pET28a (Novogen) that had been digested with *Nhe*I and *Sal*I. The ligated pET*femT* construct was transformed into *E. coli*-BL21 for protein expression and purification using metal affinity chromatography.

### Growth and Complementation assays

For growth assays, cultures of *S. aureus* were prepared by inoculating 3mL of TSB in a 13 mL polypropylene tube containing antibiotics as required and grown for 16 h. The overnight cultures were then sub-cultured into 125-mL-capacity flasks containing 25 mL TSB, or TSB modified by addition of fatty acid(s), at an initial OD_600_ of 0.01. The flasks were shaken for proper aeration, and growth (OD_600_) was monitored at hourly intervals. All cultures were grown in triplicate unless otherwise stated. Alternatively, where indicated, 200 μL cultures were inoculated into 96-well microtiter plates with an initial OD_600_ of 0.01. The growth of each culture was assessed in 4 wells and plates were incubated at 37 °C on a rotary shaker (220rpm) using a Synergy H4 Hybrid Reader (Bio Tek, Winooski, VT). Growth (OD_600_) was measured every 20 min for the indicated time.

### Reporter gene expression

Luciferase assays were conducted as previously described (Alnaseri et al., 2019; Bonn-Dunbar et al., 2025). Overnight cultures of *S. aureus* USA300 harboring appropriate reporter constructs were subcultured to an OD_600_ of 0.01 in triplicate, into a 125 mL flask containing 25mL of TSB, or TSB supplemented with 20 μM C18:2 or 50 μM C16:0 as indicated. At hourly intervals, aliquots were removed from each culture for OD_600_ measurements, and concurrently, 4×200 μL technical replicates were withdrawn from each flask for quantification of luciferase activity. For each reading, 200 μL of bacterial culture was added to individual wells of a flat-bottom, opaque white 96-well microtiter plate (Greiner Bio-One). Wells were supplemented with 20 μL of 0.1% (v/v) decanal in 40% ethanol, and luminescence was immediately measured using a BioTek Synergy H4 Hybrid Reader (integration time 1 s; gain 200). Background luminescence was corrected by subtracting the signal from the strain carrying the empty pGYlux reporter under identical conditions. Luminescence values were normalized to culture density by dividing RLU by OD_600_, generating RLU/OD_600_ measurements

### Generation of FemT antibodies

Soluble FemT were expressed in *E. coli* BL21(DE3) harboring pET-*femT* purified using Ni-NTA metal affinity chromatography as described previously (Alnaseri *et al*., 2019). Briefly, bacterial cultures were grown at 37°C in LB and supplemented with 50 µg /mL kanamycin, to an OD_600_ ∼ 0.8, before 0.1mM isopropyl thio-β-d-galactopyranoside (IPTG) was added to the culture and incubated for an additional 18 h at room temperature. Cell pellets were then resuspended in lysis buffer containing 50 mM Tris-HCl (pH 8.0), 300 mM NaCl, 10 mM imidazole, and 0.1% (v/v) Triton X-100, supplemented with EDTA-free protease inhibitor cocktail, lysozyme (1 mg/mL), DNase I (5–10 µg/mL), and MgCl_2_ (2–5 mM). Lysates were incubated on ice for an hour and clarified by centrifugation at 4,200 × *g* for 20–30 min at 4°C. The clarified supernatant was filtered through 0.45µm Acrodisc syringe filter (Pall Laboratory). The lysate was applied to 1 mL HisTrap Nickel affinity column and equilibrated with binding buffer containing 50 mM Tris-HCl (pH 8.0), 300 mM NaCl, and 10 mM imidazole. After washing the columns with wash buffer (50 mM Tris-HCl [pH 8.0], 300 mM NaCl, and 20 mM imidazole), bound His-tagged proteins were subsequently eluted using elution buffer containing (50 mM Tris-HCl [pH 8.0], 300 mM NaCl, and 250 mM imidazole). Purified proteins were dialyzed against PBS and quantified using the Bradford assay using Bio-Rad protein assay reagent. Purified proteins were submitted to ProSci Incorporated (Poway, CA) for antibody production. Polyclonal antibodies against FemT were generated in New Zealand White rabbits (n = 2 per antigen). Approximately 200 µg of purified protein was emulsified with Freund’s complete adjuvant for the primary immunization, followed by booster injections of 100 µg of protein in Freund’s incomplete adjuvant at two-week intervals for 6 weeks. Immune sera were collected 2 weeks after the final booster and stored at −80 °C until use.

### SDS-PAGE and Western Blotting

To detect FemT through western blots, USA300 was grown overnight in 3mL TSB and subcultured the next day into 25 mL TSB alone or supplemented with either C18:2 or C16:0, in 125 mL flasks. Cultures were collected for lysate preparation at indicated time points. Cells were centrifuged at 4,200 × g at 4°C for 15 min and washed with 1 × PBS. Cells were then resuspended in lysis buffer (150 mM NaCl, 50 mM Tris-HCl [pH8.0], 1% [v/v] Triton X-100, 0.5% [v/v] SDS, 0.5% sodium deoxycholate) and Pierce EDTA-free protease inhibitor cocktail, 10μg /mL lysostaphin, and I U DNase I. Lysates were incubated with for 1 h at room temperature with agitation, followed by centrifugation at 4,200 × g for 15 min at 4°C. The clarified lysate was then quantified for protein concentration using Bradford assay. For SDS-PAGE 20 µg of lysate (FemT Western blots) was mixed in 1× Laemmli sample buffer, boiled for 5 min, and resolved on 10% acrylamide gel in Tris-glycine running buffer. A prestained protein molecular weight marker (New England Biolabs) was used to estimate the sizes of protein bands. Proteins were transferred to Amersham Hybond-P FluroTrans polyvinylidene difluoride membrane (GE Healthcare) at 100 V for 1 h. Membranes were then blocked for 2 h in 5% nonfat dry milk prepared in PBS-T (1 × PBS, 0.05% Tween-20). Membranes were incubated with rabbit polyclonal anti-FemT (1:1000 dilution) overnight at 4°C, washed with PBS-T, and then incubated with IRDye 800-conjugated goat anti-rabbit secondary antibody (1:10,000 dilution) (Jackson ImmunoResearch Laboratories, Inc.) for 2 h. Blots were washed with PBS-T and imaged using an Odyssey CLx imager 32 (Li-Cor Biosciences). Images were analyzed on Fiji (ImageJ) (Schindelin *et al*., 2012).

### RNA sequencing and Read processing

Triplicate cultures of *Staphylococcus aureus* USA300 and USA300Δ*femT* were grown in tryptic soy broth (TSB) or TSB supplemented with 50 µM C16:0. 5mL of culture growing at OD_600_ ∼ 0.35–0.4, was harvested by centrifugation (4,200× g, 10min, 4°C) and pellets were resuspended in 1mL of RNA Protect (Qiagen). After incubating cells for 30 min at room temperature, cells were pelleted, and RNA was extracted using RNeasy Plus kit (Qiagen) using manufacturer’s instructions. RNA quality and integrity were assessed using the Qubit and Agilent 2100 Bioanalyzer (Agilent Technologies, USA) by the Robarts Research Institute (London, ON). Sequencing libraries were constructed and sequenced by SeqCenter, LLC (Pittsburgh, PA) to generate paired end reads. Quality-filtered reads were aligned to the *S. aureus* USA300_FPR3757 reference genome using Rsubread, and gene level counts were generated using featureCounts (Liao et al., 2014) against the corresponding gene annotation file (NC_007793.1). Differential expression analysis was performed in R using the DESeq2 package (Love et al., 2014). Genes with low read counts (<5 CPM in all samples) were excluded. Raw counts were normalized using the median ratios method, and differential expression between groups was assessed using the Wald test. Resulting *p*-values were adjusted for multiple testing using the Benjamini–Hochberg false discovery rate (FDR) method. Genes were considered significantly differentially expressed if they had an absolute log_2_ fold change ≥ 2 and an adjusted *p*-value (FDR) ≤ 0.05.

### BODIPY-vancomycin staining and confocal microscopy

*S. aureus* USA300 and USA300Δ*femT* mutant strains were cultured in TSB alone or supplemented with 50 µM C16:0 till an OD_600 ∼_ 0.35–0.40. Cells were harvested by centrifugation, washed with PBS, and fixed in 4% paraformaldehyde for 15–20 min at room temperature. Fixed cells were stained with BODIPY-vancomycin (Thermo Fisher Scientific) at concentration of 1µg/mL in PBS for 5 min, washed twice with PBS, and mounted on glass slides for imaging. A drop of antifade mounting medium (ProLong Gold Antifade Mountant, Thermofischer Scientific) was added prior to placement of coverslip (22×22 mm; VWR). Samples were visualized using a laser scanning confocal microscope (Zeiss, LSM 880) with excitation at 488 nm and emission detection between 500–550 nm. Cell volume was determined as previously described (Monteiro *et al*., 2015). Measurements of the long and short axis of the cells were performed in Fiji. Cell volume measurements were restricted to non-septated cells to ensure analysis of cells prior to septum formation. One hundred cells per strain were randomly selected from multiple fields of view. The volume was calculated using the formula: 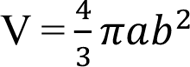, where V is the volume, a and b are radii for long and the short axis respectively.

### Cell respiration assay

Respiration was measured using 5-cyano-2,3-ditolyl tetrazolium chloride (CTC) stain as previously described for *S. aureus* (Lewis *et al*., 2015). USA300 pALC, USA300Δ*femT* pALC, and the complemented strain USA300Δ*femT* pALC*femT* were grown in 25 mL TSB or TSB supplemented with 50 µM C16:0 and grown to an OD_600_ of 0.3–0.4. Next, 2 mL of each culture was pelleted by centrifugation at 4,200 × g at room temperature, washed once with 1 mL of 1× PBS, and resuspended in 650 µL of PBS containing 4.5 mM CTC. Triplicate aliquots (200 µL) of each stained suspension were transferred to a black-walled 96-well plate (Agilent). Fluorescence was measured using a microplate reader (BioTek Synergy) with excitation/emission settings of ∼485/590 nm. Fluorescence values (RFU) were normalized to cell density (OD_600_). Heat-killed controls were prepared by incubating cell suspensions at 95 °C for 10 min prior to CTC addition.

### Membrane Fluidity assay

Membrane fluidity was assessed using Laurdan generalized polarization (GP) as described previously (Wenzel *et al*., 2018). Overnight cultures of USA300 and USA300Δ*femT* were diluted to a final OD_600_ of 0.01 into 25 mL of TSB or TSB supplemented with either 50 µM C16:0 or 50 µM C18:1 in 125-mL flasks. Cultures were grown in biological triplicate at 37°C with shaking until mid-exponential phase. Cells were harvested, washed with Laurdan buffer, and resuspended in Laurdan buffer (137 mM NaCl, 2.7 mM KCl, 10 mM Na_2_HPO_4_, 1.8 mM KH_2_PO_4_, 0.2% glucose, and 1% DMSO) containing Laurdan (Cayman) at a final concentration of 10 µM. Cultures treated with 100 µM linoleic acid for 15-20 min at 37°C prior to Laurdan staining served as a membrane fluidization control. Cell suspensions were normalized to an OD_600_ of 0.4 and incubated for 30 min at 37°C in the dark. Following incubation, cells were washed with Laurdan buffer and 4 × 200µl technical replicates were transferred to black wall 96-well plate (Agilent) for fluorescence measurement using microplate reader (BioTek Synergy) with excitation at 350 nm, and emission recorded at 435 nm and 500 nm. Generalized polarization was calculated as GP = (I435-I500)/ (I435+I500).

### Membrane potential quantification

Membrane potential was measured using DiOC_2_(3) (3,3′-diethyloxacarbocyanine iodide; Thermo Fisher Scientific) as described previously (Dombach *et al*., 2024). Overnight cultures of USA300 and USA300Δ*femT* were diluted into 25 mL TSB or TSB supplemented with 50 µM C16:0 and grown in 125-mL flasks (37°C, 220 rpm) to mid-exponential phase in biological triplicate. Cells were harvested by centrifugation (5,000 × g, 5 min), washed once with PBS, and resuspended in PBS to an OD_600_ of 0.4. DiOC_2_(3) was added to a final concentration of 30 µM, and samples were incubated at 37°C for 30 min in the dark. Cells were washed once with PBS and resuspended in the same buffer. Aliquots (200 µL) were dispensed into four technical replicate wells per biological replicate in a black 96-well plate. Fluorescence was measured using a microplate reader (BioTek Synergy) with excitation at 488 nm and emission recorded at 530 nm (green) and 610 nm (red). Membrane potential was expressed as the ratio of red to green fluorescence. Where indicated, cells treated with a final concentration of 1 μg/mL gramicidin (Sigma-Aldrich) for 5 minutes and were used as a depolarized control.

### Intracellular ROS quantification

Intracellular reactive oxygen species were measured using 2′,7′-dichlorodihydrofluorescein diacetate (DCFH-DA,Sigma) as described previously (Wang and Joseph, 1999). Cultures were grown in tryptic soy broth supplemented with 50 µM C16:0 to early-exponential phase (OD_600_∼ 0.25). DCFH-DA was added to a final concentration of 20 µM, and cultures were incubated in the dark for 30 min at 37°C. Hydrogen peroxide-treated cultures served as a positive control. Exponential-phase cultures were exposed to 1 mM H_2_O_2_ for 30 min at 37°C prior to DCFH-DA staining. Following incubation, cells were washed once with PBS to remove excess dye and resuspended in PBS. Samples were transferred to a black 96-well plate, and fluorescence was measured at excitation and emission wavelengths of 485 and 520 nm, respectively. Background-corrected fluorescence was normalized to OD_600_ and reported as relative fluorescence units per OD_600._

### Metabolomics

Overnight cultures of *S. aureus* USA300 and USA300Δ*femT* were used to inoculate 25 mL TSB at an initial OD600 of ∼0.01 with or without 50 µM C16:0 and grown till OD_600_ ∼ 0.35–0.4. Next, 10 mL of culture was harvested by centrifugation (3,000 × g, 10 min, 4°C), washed with ice-cold 1× PBS, and centrifuged again (3,000 × g, 5 min, 4°C). Pellets were resuspended in ice cold acetonitrile:methanol:water (40:40:20, v/v/v) containing 0.1 M formic acid and transferred to lysing matrix tubes (MP Biomedicals) compatible with the FastPrep-24 instrument (MP Biomedicals). Cell lysis was performed by bead beating using a FastPrep-24 instrument for two cycles of 60 s at 6.0 m s⁻¹, with cooling on ice between cycles. Lysates were clarified by centrifugation (15 min, 4°C), transferred to fresh tubes, centrifuged again (10 min), and filtered through 0.22 µm filters. Filtrates were stored at −80°C until analysis.

Metabolomic profiling was performed by the University of Calgary Centre for Metabolomics and Mass Spectrometry Research Facility (CMRF) using methodology as described previously (Groves et al., 2022; Mager et al., 2020; Rydzak et al., 2022). Analysis was performed on Q Exactive™ Hybrid Quadrupole-Orbitrap™ Mass Spectrometer (Thermo-Fisher) coupled to a Vanquish™ UHPLC System (Thermo-Fisher). Briefly, metabolites in the samples were resolved with a Syncronis^TM^ Hydrophilic interaction Liquid Chromatography (HILIC) column (2.1mm x 100mm x 1.7µm, Thermo-Fisher) on a Vanquish^TM^ Ultra-High-Performance Liquid Chromatography (UHPLC) platform (ThermoScientific) using 15 min two-solvent gradient method (20mM ammonium formate at pH 3.0 in HPLC-grade water and HPLC-grade acetonitrile with 0.1% formic acid). The mass spectrometer was run at a resolution of 140,000 scanning from 50-750m/z. Metabolite data was analyzed by El-MAVEN software package (Clasquin et al., 2012; Melamud et al., 2010). Metabolites were identified by matching observed m/z signals (+/-10ppm) and chromatographic retention times to those observed from commercial metabolite standards (MSMLS**^TM^**Sigma-Aldrich). Next, metabolites were quantified by comparison to an eight-point quantification curve of metabolite standards. Metabolites were ranked by adjusted *p*-values following ANOVA with Benjamini–Hochberg FDR correction. Heatmaps were generated using row-wise z-score–normalized intensities, with hierarchical clustering based on Euclidean distance and Ward’s linkage.

### Lipidomics

Overnight cultures of *S. aureus* USA300 and the USA300*ΔfemT* mutant were diluted into TSB and grown to OD_600_ ∼ 0.35–0.4. Cultures were then exposed to 500 µM C18:1 and grown to OD _600_ ∼ 1.0, after which cells were harvested by centrifugation at 5,000 × g for 10 min at 4 °C, washed with cold PBS, and cell pellets were flash-frozen in liquid nitrogen. Samples (n = 4 biological replicates per condition) were analyzed at the University of Calgary Central Metabolomics and Lipidomics Research Facility (CMRF). Lipid extraction was performed using a modified Folch liquid–liquid extraction protocol (Folch et al., 1951) employing dichloromethane and methanol, with 0.2 M KCl used for phase separation. Solvent volumes were adjusted to accommodate low sample amounts (Zardini Buzatto, Kwon and Li, 2020). Samples were randomized during preparation and analysis to minimize batch effects. Aliquots of 2.0 × 10⁹ CFU were sequentially mixed with 5.2 µL internal standard mixture (SPLASH Lipidomix, Avanti Polar Lipids), 261.0 µL methanol, and 533.0 µL dichloromethane, followed by homogenization for 60 s. Samples were centrifuged briefly and vortexed for 30 s with 198.0 µL of 0.2 M KCl, equilibrated for 10 min at 4 °C, and centrifuged for 10 min at 12,000 rpm at 4 °C. The lower organic phase was collected and evaporated to dryness using a SpeedVac concentrator. Remaining organic layers were pooled, vortexed, and split into equal-volume aliquots for quality control (QC), which were also dried. Dried extracts were stored at −80 °C, protected from light, for up to 24 h. Lipid extracts were analyzed by reversed-phase liquid chromatography coupled to high-resolution mass spectrometry (LC–MS/MS) using a Thermo Vanquish Horizon UHPLC system equipped with a Waters Acquity Premier CSH C18 column (1.7 µm, 2.1 × 100 mm) coupled to a Thermo Q-Exactive HF Orbitrap mass spectrometer. Samples were analyzed in both positive and negative ionization modes, and mass spectra were acquired over an m/z range of 140–2000 using data-dependent acquisition. Mobile phases consisted of 10 mM ammonium formate in 2:2:1 methanol/acetonitrile/water (mobile phase A) and 10 mM ammonium formate in 96:3:2 2-propanol/acetonitrile/water (mobile phase B), and lipids were separated using a 16.5-min gradient elution. Chromatograms were processed using LipidQuest (Buzatto Research Group), which performs retention time correction, peak picking, alignment, polarity merging, and lipid annotation. Lipid identities were assigned by matching accurate mass and MS/MS fragmentation spectra to public databases including LIPID MAPS and LipidBlast, and when available to authentic standards. Peak intensities were normalized to the most structurally similar deuterated internal standard. A commercially available 10-hydroxystearic acid (10-HSA; Ambeed) standard was used to support metabolite identification. Retention time and mass spectral features of the standard were used to aid annotation of 10-HSA in samples. Statistical analyses were performed in Python using normalized lipid intensities. Lipidome differences were evaluated using t-tests with Benjamini–Hochberg for false discovery rate correction. Graphs were generated using normalized lipid abundance values derived from the processed LC–MS/MS peak tables provided by CMRF.

### Subcutaneous infection model

Mouse experiments were carried out following the Canadian Council on Animal Care Guide to the Care and Use of Experimental Animals and the animal protocol was approved by the Animal Use Subcommittee at the University of Western Ontario (Protocol 2024-082). Eight- to twelve-week-old male and female C57BL/6 mice were purchased from Jackson Laboratory (USA). Overnight cultures of *S. aureus* were diluted 1:50 into fresh tryptic soy broth (TSB) and grown at 37 °C with shaking till an OD_600_ ∼ 3. Bacterial cells were harvested by centrifugation, washed, and resuspended in sterile Hanks’ balanced salt solution (HBSS) to the desired inoculum. Mice were anesthetized with isoflurane, and approximately 5 × 10⁷ CFU of bacteria in 50 µL HBSS were injected subcutaneously into the shaved lower flank. Infected animals were monitored daily for body weight, general health, and lesion development for up to 3 days. On day 3 post-infection, mice were euthanized, and lesion size was measured using a caliper. Lesions were surgically excised, homogenized twice for 5 min in 4 mL HBSS using metal beads in 5 mL screw-cap tubes, and bacterial burdens were determined by serial dilution and plating on mannitol salt agar (MSA).

### Data analyses

All data visualization and statistical analysis were performed using GraphPad Prism (version 10.4.1). Unless specified otherwise, experiments were conducted with triplicate cultures, and data are presented as the mean ± standard error of the mean (SEM). Statistical significance, defined in the figure legends, was determined using either a one-way or two-way ANOVA with multiple comparisons, as appropriate for the experimental design.

## ACKNOWLEDGMENTS

This work was supported by Canadian Institutes of Health Research grant PJT-463397 (M.J.M.). A.T was a recipient of an R.G.E. Murray Graduate Scholarship award from Western University, Department of Microbiology and Immunology. We thank the Calgary Metabolomics Research Facility (CMRF) for performing lipidomics (Dr. Adriana Zardini Buzatto) and metabolomics analyses (Dr. Annegret Ulke-Lemée) for this project. We also acknowledge the IMPAKT Facility for access to confocal microscopy instrumentation. Figure for graphical abstract was created using Biorender

## REFERENCES

Adams, O. et al. (2021) “Cryo-EM structure and resistance landscape of M. tuberculosis MmpL3: An emergent therapeutic target,” Structure, 29(10), pp. 1182–1191.e4.

Adebusuyi, A.A. and Foght, J.M. (2011) “An alternative physiological role for the EmhABC efflux pump in Pseudomonas fluorescens cLP6a,” BMC microbiology, 11.

Alenazy, R. (2022) “Drug Efflux Pump Inhibitors: A Promising Approach to Counter Multidrug Resistance in Gram-Negative Pathogens by Targeting AcrB Protein from AcrAB-TolC Multidrug Efflux Pump from Escherichia coli,” Biology.

Alnaseri, H. et al. (2015) “Inducible Expression of a Resistance-Nodulation-Division-Type Efflux Pump in Staphylococcus aureus Provides Resistance to Linoleic and Arachidonic Acids,” Journal of Bacteriology. 2015/03/23, 197(11), pp. 1893–1905.

Alnaseri, H. et al. (2019) “DNA binding and sensor specificity of FarR, a novel tetr family regulator required for induction of the fatty acid efflux pump FarE in staphylococcus aureus,” Journal of Bacteriology, 201(3).

Arsic, B. et al. (2012) “Induction of the staphylococcal proteolytic cascade by antimicrobial fatty acids in community acquired methicillin resistant Staphylococcus aureus,” PloS one, 7(9).

Bae, T. and Schneewind, O. (2006) “Allelic replacement in Staphylococcus aureus with inducible counter-selection,” Plasmid, 55(1), pp. 58–63.

Balibar, C.J. et al. (2010) “cwrA, a gene that specifically responds to cell wall damage in Staphylococcus aureus,” Microbiology, 156(5), pp. 1372–1383.

Balibar, C.J., Shen, X. and Tao, J. (2008) “The Mevalonate Pathway of Staphylococcus aureus,” Journal of Bacteriology, 191(3), p. 851.

Beetham, C.M. et al. (2024) “Histidine transport is essential for the growth of Staphylococcus aureus at low pH,” PLOS Pathogens, 20(1), p. e1011927.

Bender, R.A. (2012) “Regulation of the Histidine Utilization (Hut) System in Bacteria,” Microbiology and Molecular Biology Reviews, 76(3), pp. 565–584.

Bonn, C.M. et al. (2023) “Repeated Emergence of Variant TetR Family Regulator, FarR, and Increased Resistance to Antimicrobial Unsaturated Fatty Acid among Clonal Complex 5 Methicillin-Resistant Staphylococcus aureus,” Antimicrobial Agents and Chemotherapy, 67(3).

Bonn-Dunbar, C.M. et al. (2025) “Widespread emergence of Staphylococcus aureus with variant FarR regulators and enhanced resistance to antimicrobial fatty acids within clonal complex CC5, CC8, and CC97 strains from human and bovine hosts,” Microbiology Spectrum, 13(12), pp. e02278–25.

Budin, I. et al. (2018) “Viscous control of cellular respiration by membrane lipid composition,” Science, 362(6419), pp. 1186–1189.

Burckhardt, R.M., Buckner, B.A. and Escalante-Semerena, J.C. (2019) “Staphylococcus aureus modulates the activity of acetyl-Coenzyme A synthetase (Acs) by sirtuin-dependent reversible lysine acetylation,” Molecular microbiology, 112(2), pp. 588–604.

Chatterjee, I. et al. (2011a) “Staphylococcus aureus ClpC is involved in protection of carbon-metabolizing enzymes from carbonylation during stationary growth phase,” International Journal of Medical Microbiology, 301(4), pp. 341–346.

Chatterjee, I. et al. (2011b) “Staphylococcus aureus ClpC is involved in protection of carbon-metabolizing enzymes from carbonylation during stationary growth phase,” International journal of medical microbiology: IJMM, 301(4), pp. 341–346.

Chen, Y. et al. (2022) “Transcriptomic Responses and Survival Mechanisms of Staphylococci to the Antimicrobial Skin Lipid Sphingosine,” Antimicrobial Agents and Chemotherapy, 66(2), pp. e00569–21.

Chen, Z. et al. (2018) “Acetylome Profiling Reveals Extensive Lysine Acetylation of the Fatty Acid Metabolism Pathway in the Diatom Phaeodactylum tricornutum,” Molecular & cellular proteomics: MCP, 17(3), pp. 399–412.

Clasquin, M.F., Melamud, E. and Rabinowitz, J.D. (2012) “LC-MS data processing with MAVEN: a metabolomic analysis and visualization engine,” *Current protocols in bioinformatics*, Chapter 14(SUPPL.37).

Collins, M.D. and Jones, D. (1981) “Distribution of isoprenoid quinone structural types in bacteria and their taxonomic implication,” Microbiological Reviews, 45(2), pp. 316–354.

Corrigan, R.M. and Foster, T.J. (2009) “An improved tetracycline-inducible expression vector for Staphylococcus aureus,” Plasmid, 61(2), pp. 126–129.

Crosby, H.A. et al. (2012) “System-wide studies of N-lysine acetylation in Rhodopseudomonas palustris reveal substrate specificity of protein acetyltransferases,” The Journal of biological chemistry, 287(19), pp. 15590–15601.

Desai, J. et al. (2016) “Structure, Function, and Inhibition of Staphylococcus aureus Heptaprenyl Diphosphate Synthase,” ChemMedChem, 11(17), pp. 1915–1923.

Ericson, M.E. et al. (2017) “Role of fatty acid kinase in cellular lipid homeostasis and SaeRS-dependent virulence factor expression in Staphylococcus aureus,” mBio, 8(4).

Folch, J. et al. (1951) “PREPARATION OF LIPIDE EXTRACTS FROM BRAIN TISSUE*,” Journal of Biological Chemistry, 191, pp. 833–841.

Frank, M.W. et al. (2020) “Host Fatty Acid Utilization by Staphylococcus aureus at the Infection Site,” mBio, 11(3).

Fraud, S. et al. (2008) “MexCD-OprJ multidrug efflux system of Pseudomonas aeruginosa: Involvement in chlorhexidine resistance and induction by membrane-damaging agents dependent upon the AlgU stress response sigma factor,” Antimicrobial Agents and Chemotherapy, 52(12), pp. 4478–4482.

Götz, F. and Mayer, S. (2013) “Both terminal oxidases contribute to fitness and virulence during organ-specific Staphylococcus aureus colonization,” mBio, 4(6).

Groves, R.A. et al. (2022) “Methods for Quantifying the Metabolic Boundary Fluxes of Cell Cultures in Large Cohorts by High-Resolution Hydrophilic Liquid Chromatography Mass Spectrometry,” Analytical Chemistry, 94(25), pp. 8874–8882.

Holm, L. (2020) “Using Dali for Protein Structure Comparison BT - Structural Bioinformatics: Methods and Protocols,” in Z. Gáspári (ed.). New York, NY: Springer US, pp. 29–42.

Horton, R.M. et al. (1993) “Gene splicing by overlap extension,” Methods in Enzymology, 217(C), pp. 270–279.

Huang, L. et al. (2022) “Molecular Basis of Rhodomyrtone Resistance in Staphylococcus aureus,” mBio, 13(1

Jerga, A. et al. (2007) “Identification of a Soluble Diacylglycerol Kinase Required for Lipoteichoic Acid Production in Bacillus subtilis,” Journal of Biological Chemistry, 282(30), pp. 21738–21745.

Kearns, D.B. et al. (2004) “Genes governing swarming in Bacillus subtilis and evidence for a phase variation mechanism controlling surface motility,” Molecular Microbiology, 52(2), pp. 357–369.

Kénanian, G. et al. (2019a) “Permissive Fatty Acid Incorporation Promotes Staphylococcal Adaptation to FASII Antibiotics in Host Environments,” Cell reports, 29(12), pp. 3974–3982.e4.

Kénanian, G. et al. (2019b) “Permissive Fatty Acid Incorporation Promotes Staphylococcal Adaptation to FASII Antibiotics in Host Environments,” Cell Reports, 29(12), pp. 3974–3982.e4.

Kenny, J.G. et al. (2009) “The Staphylococcus aureus Response to Unsaturated Long Chain Free Fatty Acids: Survival Mechanisms and Virulence Implications,” PLOS ONE, 4(2), p. e4344.

Kinkel, T.L. et al. (2013) “The Staphylococcus aureus SrrAB two-component system promotes resistance to nitrosative stress and hypoxia,” mBio, 4(6).

Kuiack, R.C. et al. (2023) “The fadXDEBA locus of Staphylococcus aureus is required for metabolism of exogenous palmitic acid and in vivo growth,” Molecular Microbiology, 120(3), pp. 425–438.

Larosa, V. and Remacle, C. (2018) “Insights into the respiratory chain and oxidative stress,” Bioscience Reports, 38(5), p. BSR20171492.

Lemire, J. et al. (2017) “Metabolic defence against oxidative stress: the road less travelled so far,” Journal of Applied Microbiology, 123(4), pp. 798–809.

Lewis, A.M. et al. (2015) “Examination of the Staphylococcus aureus Nitric Oxide Reductase (saNOR) Reveals its Contribution to Modulating Intracellular NO Levels and Cellular Respiration,” Molecular microbiology, 96(3), p. 651.

Liao, Y., Smyth, G.K. and Shi, W. (2014) “featureCounts: an efficient general purpose program for assigning sequence reads to genomic features,” Bioinformatics, 30(7), pp. 923–930.

Liechti, G.W. and Goldberg, J.B. (2026) “Glutamate metabolism: an essential component for persistence in Staphylococcus aureus,” Microbiology Spectrum [Preprint].

Lithgow, J.K., Ingham, E. and Foster, S.J. (2004) “Role of the *hprT-ftsH* locus in Staphylococcus aureus,” Microbiology, 150(2), pp. 373–381

Liu, G.Y. et al. (2005) “Staphylococcus aureus golden pigment impairs neutrophil killing and promotes virulence through its antioxidant activity,” The Journal of experimental medicine, 202(2), pp. 209–215.

Liu, Y. et al. (2022) “Metabolic Mechanism and Physiological Role of Glycerol 3-Phosphate in Pseudomonas aeruginosa PAO1,” mBio, 13(6).

Lo, R. et al. (2009) “Cystathionine γ-Lyase is a component of cystine-mediated oxidative defense in lactobacillus reuteri br11,” Journal of Bacteriology, 191(6), pp. 1827–1837.

Van Loi, V. et al. (2023) “Staphylococcus aureus adapts to the immunometabolite itaconic acid by inducing acid and oxidative stress responses including S-bacillithiolations and S-itaconations,” Free Radical Biology and Medicine, 208, pp. 859–876.

Lopez, M.S. et al. (2017a) “Host-derived fatty acids activate type VII secretion in Staphylococcus aureus,” Proceedings of the National Academy of Sciences of the United States of America, 114(42), pp. 11223–11228.

Lopez, M.S. et al. (2017b) “Host-derived fatty acids activate type VII secretion in Staphylococcus aureus,” Proceedings of the National Academy of Sciences of the United States of America, 114(42), pp. 11223–11228.

Love, M.I., Huber, W. and Anders, S. (2014) “Moderated estimation of fold change and dispersion for RNA-seq data with DESeq2,” Genome Biology, 15(12), pp. 550-.

Lu, Y. et al. (2023) “Modulation of MRSA virulence gene expression by the wall teichoic acid enzyme TarO,” Nature Communications, 14(1).

Lund, P.A. (2013) “Bacterial Stress Responses,” pp. 3–22.

Ma, D. et al. (1995) “Genes acrA and acrB encode a stress-induced efflux system of Escherichia coli,” Molecular microbiology, 16(1), pp. 45–55

Mager, L.F. et al. (2020) “Microbiome-derived inosine modulates response to checkpoint inhibitor immunotherapy,” *Science (New York*, N.Y.), 369(6510), pp. 1481–1489.

Melamud, E., Vastag, L. and Rabinowitz, J.D. (2010) “Metabolomic analysis and visualization engine for LC-MS data,” Analytical chemistry, 82(23), pp. 9818–9826.

Mesak, L.R., Yim, G. and Davies, J. (2009) “Improved lux reporters for use in Staphylococcus aureus,” Plasmid, 61(3), pp. 182–187.

Misic, A.M. et al. (2016) “ Divergent Isoprenoid Biosynthesis Pathways in Staphylococcus Species Constitute a Drug Target for Treating Infections in Companion Animals,” mSphere, 1(5).

Monteiro, J.M. et al. (2015) “Cell shape dynamics during the staphylococcal cell cycle,” Nature Communications 2015 6:1, 6(1), pp. 8055-.

Nakamura, H., Hachiya, N. and Tojo, T. (1978) “Second Acriflavine Sensitivity Mutation, acrB, in Escherichia coli K-12,” Journal of Bacteriology, 134(3), pp. 1184–1187.

Nakayasu, E.S. et al. (2017) “Ancient regulatory role of lysine acetylation in central metabolism,” mBio, 8(6).

Nguyen, M.T. et al. (2019) “Inactivation of farR Causes High Rhodomyrtone Resistance and Increased Pathogenicity in Staphylococcus aureus,” Frontiers in microbiology, 10(MAY).

Novick, R.P. (1991) “Genetic systems in staphylococci,” Methods in enzymology, 204(C), pp. 587–636.

Parsons, J.B. et al. (2014) “Incorporation of extracellular fatty acids by a fatty acid kinase-dependent pathway in Staphylococcus aureus,” Molecular microbiology, 92(2), pp. 234– 245.

Pathania, A. et al. (2021) “(P)ppgpp/gtp and malonyl-coa modulate staphylococcus aureus adaptation to fasII antibiotics and provide a basis for synergistic bi-therapy,” mBio, 12(1), pp. 1–15.

Pelz, A. et al. (2005) “Structure and biosynthesis of staphyloxanthin from Staphylococcus aureus,” Journal of Biological Chemistry, 280(37), pp. 32493–32498.

Piewngam, P. and Otto, M. (2024) “Staphylococcus aureus colonisation and strategies for decolonisation,” The Lancet Microbe, 5(6), pp. e606–e618.

Planet, P.J. et al. (2015) “Parallel Epidemics of Community-Associated Methicillin-Resistant Staphylococcus aureus USA300 Infection in North and South America,” The Journal of Infectious Diseases, 212(12), pp. 1874–1882.

Poole, R.K. and Cook, G.M. (2000) “Redundancy of aerobic respiratory chains in bacteria? Routes, reasons and regulation,” Advances in Microbial Physiology, 43, pp. 165–224.

Quiblier, C. et al. (2011) “Contribution of SecDF to Staphylococcus aureus resistance and expression of virulence factors,” BMC Microbiology, 11, p. 72.

Quiblier, C. et al. (2013) “The Staphylococcus aureus membrane protein SA2056 interacts with peptidoglycan synthesis enzymes,” Antibiotics, 2(1), pp. 11–27.

Radka, C.D., Batte, J.L., Frank, M.W., Rosch, J.W., et al. (2021) “Oleate hydratase (OhyA) is a virulence determinant in Staphylococcus aureus,” Microbiology Spectrum, 9(3), pp. e01546–21.

Radka, C.D., Batte, J.L., Frank, M.W., Young, B.M., et al. (2021) “Structure and mechanism of Staphylococcus aureus oleate hydratase (OhyA),” Journal of Biological Chemistry, 296.

Rizo, J. and Encarnación-Guevara, S. (2024) “Bacterial protein acetylation: mechanisms, functions, and methods for study,” Frontiers in Cellular and Infection Microbiology, 14, p. 1408947.

Rohrer, S. et al. (1999) “The essential Staphylococcus aureus gene fmhB is involved in the first step of peptidoglycan pentaglycine interpeptide formation,” Proceedings of the National Academy of Sciences, 96(16), pp. 9351–9356.

Rydzak, T. et al. (2022) “Metabolic preference assay for rapid diagnosis of bloodstream infections,” Nature Communications 2022 13:1, 13(1), pp. 2332-.

Santajit, S. and Indrawattana, N. (2016) “Mechanisms of Antimicrobial Resistance in ESKAPE Pathogens,” BioMed research international. 2016/05/05, 2016, p. 2475067.

Sati, H. et al. (2025) “The WHO Bacterial Priority Pathogens List 2024: a prioritisation study to guide research, development, and public health strategies against antimicrobial resistance,” The Lancet Infectious Diseases, 25(9), pp. 1033–1043.

Schindelin, J. et al. (2012) “Fiji: an open-source platform for biological-image analysis,” Nature methods, 9(7), pp. 676–682.

Schon, E.A. (2018) “Bioenergetics through thick and thin Membrane fluidity influences the efficiency of oxidative energy metabolism,” Science, 362(6419), pp. 1114–1115.

Somerville, G.A. et al. (2002) “Staphylococcus aureus aconitase inactivation unexpectedly inhibits post-exponential-phase growth and enhances stationary-phase survival,” Infection and Immunity, 70(11), pp. 6373–6382.

Soutourina, O. et al. (2010) “The Pleiotropic CymR Regulator of Staphylococcus aureus Plays an Important Role in Virulence and Stress Response,” PLOS Pathogens, 6(5), p. e1000894.

Starai, V.J. and Escalante-Semerena, J.C. (2004) “Identification of the protein acetyltransferase (Pat) enzyme that acetylates acetyl-CoA synthetase in Salmonella enterica,” Journal of Molecular Biology, 340(5), pp. 1005–1012.

Su, C.-C. et al. (2019) “MmpL3 is a lipid transporter that binds trehalose monomycolate and phosphatidylethanolamine,” Proceedings of the National Academy of Sciences, 116(23), pp. 11241–11246.

Subramanian, C. et al. (2019) “Oleate hydratase from Staphylococcus aureus protects against palmitoleic acid, the major antimicrobial fatty acid produced by mammalian skin,” Journal of Biological Chemistry, 294(23), pp. 9285–9294.

Thacharodi, A. et al. (2025) “Methicillin-resistant Staphylococcus aureus is raising global concern as it overcomes immune challenges through various virulence mechanisms,” iScience, 29(1).

Thurlow, L.R., Joshi, G.S. and Richardson, A.R. (2012) “Virulence Strategies of the Dominant USA300 Lineage of Community Associated Methicillin Resistant Staphylococcus aureus (CA-MRSA),” FEMS immunology and medical microbiology, 65(1), p. 5.

Tsuge, K., Ohata, Y. and Shoda, M. (2001) “Gene yerP, Involved in Surfactin Self-Resistance in Bacillus subtilis,” Antimicrobial Agents and Chemotherapy, 45(12), p. 3566.

Varadi, M. et al. (2022) “AlphaFold Protein Structure Database: massively expanding the structural coverage of protein-sequence space with high-accuracy models,” Nucleic Acids Research, 50(D1), pp. D439–D444.

Whittle, E.E. et al. (2024) “Efflux pumps mediate changes to fundamental bacterial physiology via membrane potential,” mBio, 15(10).

Yamasaki, S. et al. (2023) “Drug resistance and physiological roles of RND multidrug efflux pumps in Salmonella enterica, Escherichia coli and Pseudomonas aeruginosa,” *Microbiology (Reading*, England), 169(6).

Zardini Buzatto, A., Kwon, B.K. and Li, L. (2020) “Development of a NanoLC-MS workflow for high-sensitivity global lipidomic analysis,” Analytica chimica acta, 1139, pp. 88–99.

Zhao, L. et al. (2024) “Inhibitor binding and disruption of coupled motions in MmpL3 protein: Unraveling the mechanism of trehalose monomycolate transport,” Protein Science: A Publication of the Protein Society, 33(10), pp. e5166–e5166.

Zhong, Y. et al. (2025) “Design, synthesis and optimization of TarO inhibitors as multifunctional antibiotics against Methicillin-resistant Staphylococcus aureus,” npj Antimicrobials and Resistance, 3(1), p. 28.

