## Supplemental Figures for "The RND family efflux pump FemT contributes to lipid homeostasis in *Staphylococcus aureus”*"

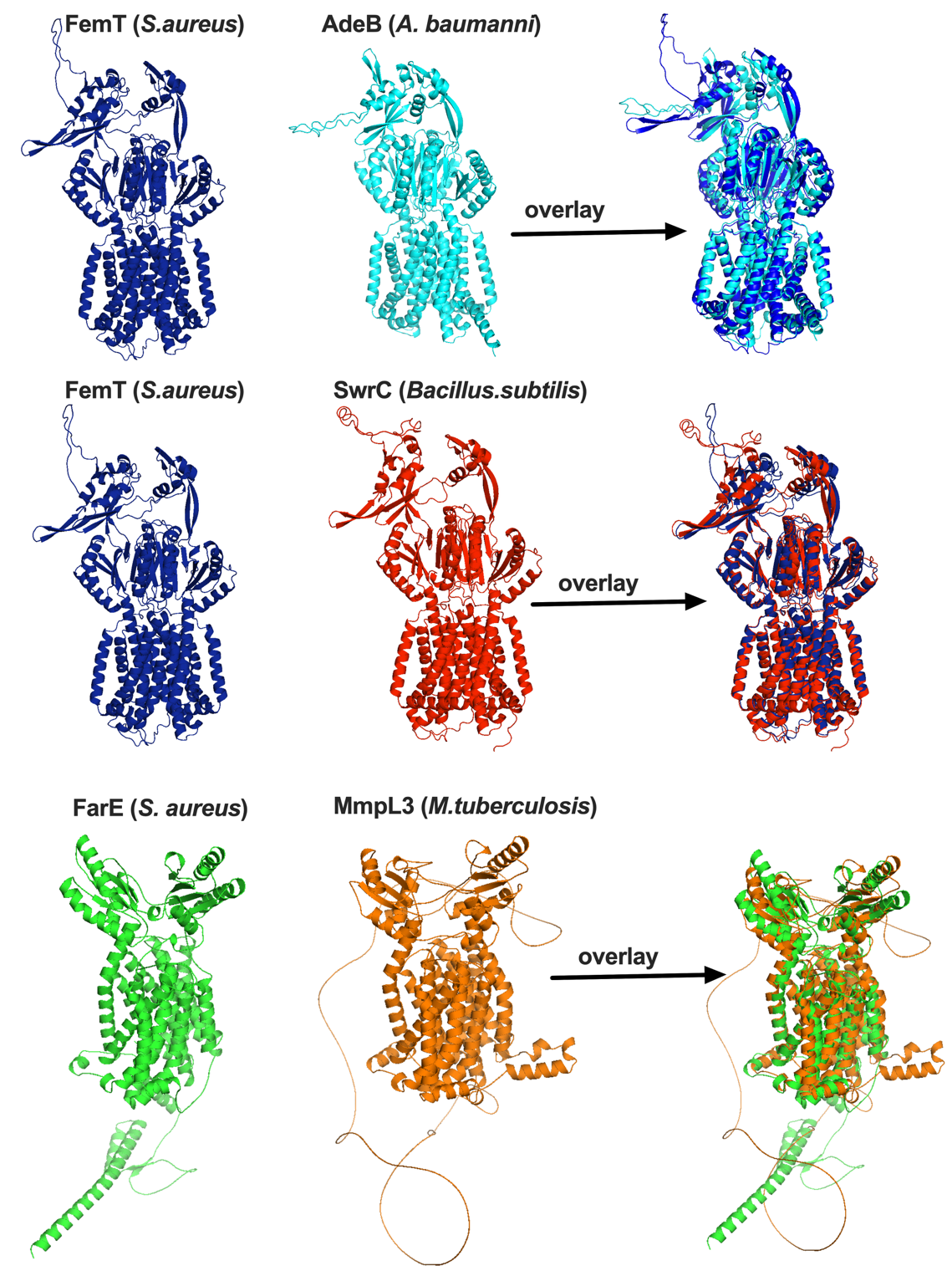

**Supplementary Figure 1.** Structural comparison of FemT and FarE with representative transporters. Predicted structures of *Staphylococcus aureus* FemT and FarE were generated using AlphaFold. FemT is shown alongside SwrC from *Bacillus subtilis* and AdeB from *Acinetobacter baumannii*, while FarE is shown alongside MmpL3 from *Mycobacterium tuberculosis*. Individual protein structures are displayed in the left and middle panels, and structural overlays following alignment are shown in the right panels for each comparison. Proteins are colored as follows: FemT (blue), AdeB (cyan), SwrC (red), FarE (green), and MmpL3 (orange). Images were generated and aligned using PyMOL.

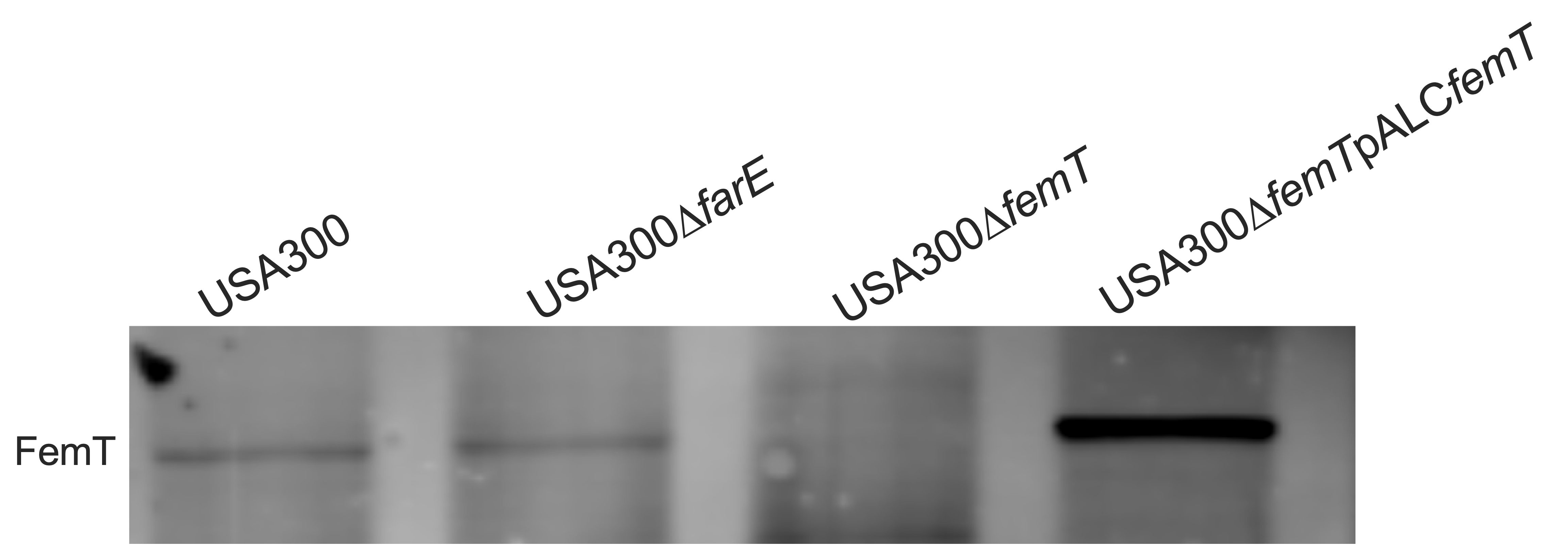

**Supplementary Figure 2.** Detection of FemT protein expression in USA300 and derivative strains. Western blot analysis of FemT in whole-cell lysates from USA300, USA300Δ*farE*, USA300Δ*femT*, and USA300Δ*femT* complemented with pALC*femT*. Equal amounts of protein (20µg) were resolved by SDS-PAGE and probed with anti-FemT antibodies. FemT is detected in USA300 and USA300Δ*farE*, absent in USA300Δ*femT*, and restored in the complemented strain.

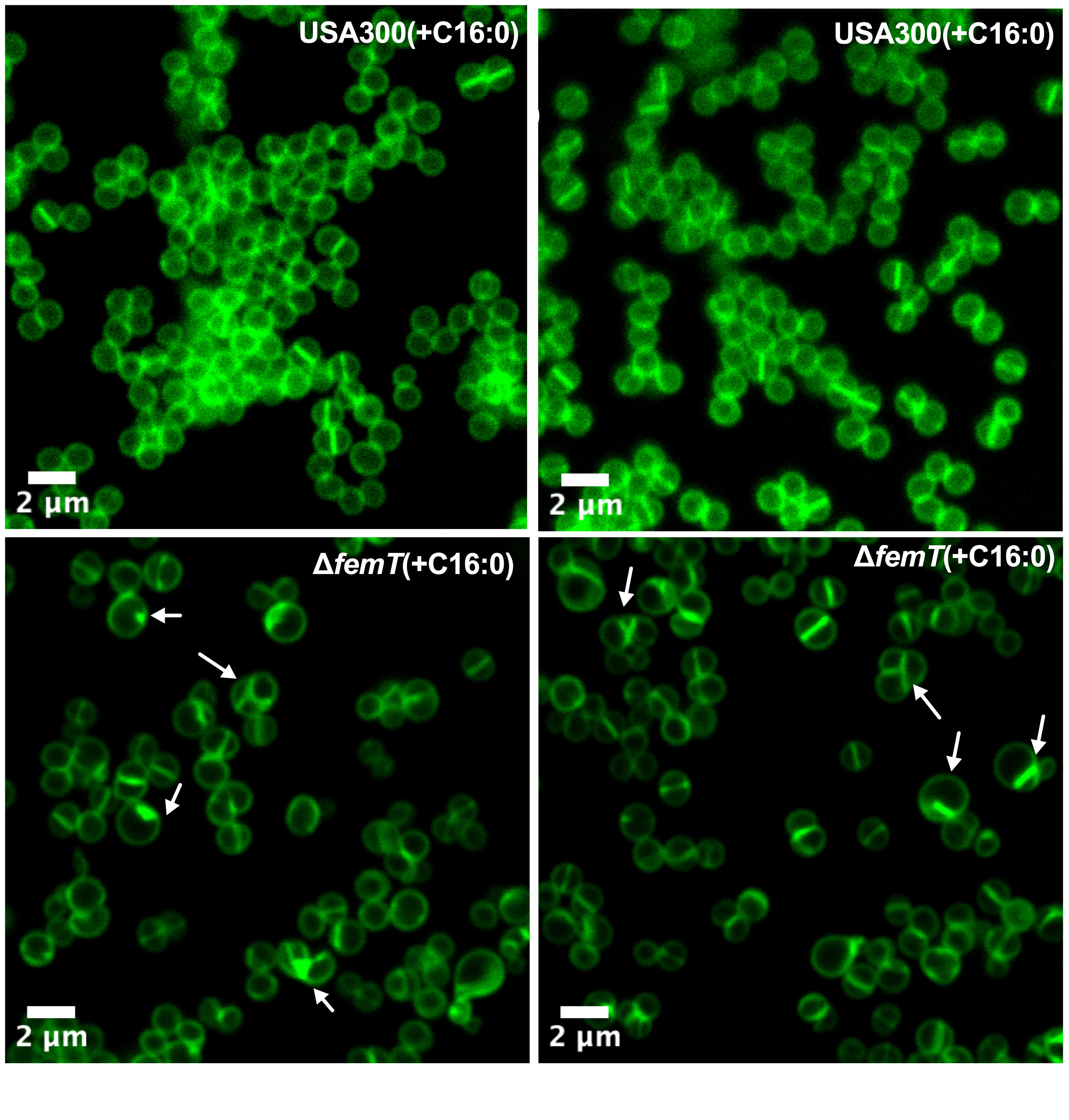
**Supplementary Figure 3: Representative images of USA300∆*femT* grown in palmitic acid**. Cells were grown in TSB supplemented with 50 µM C16:0, harvested at mid-log phase (OD_600_~0.3–0.4), washed, and stained with BODIPY-vancomycin to visualize the cell envelope. Representative fields are shown in panels A and B. Arrows indicate cells exhibiting aberrant septal staining patterns and increased fluorescence intensity. Scale bars, 2 µm.

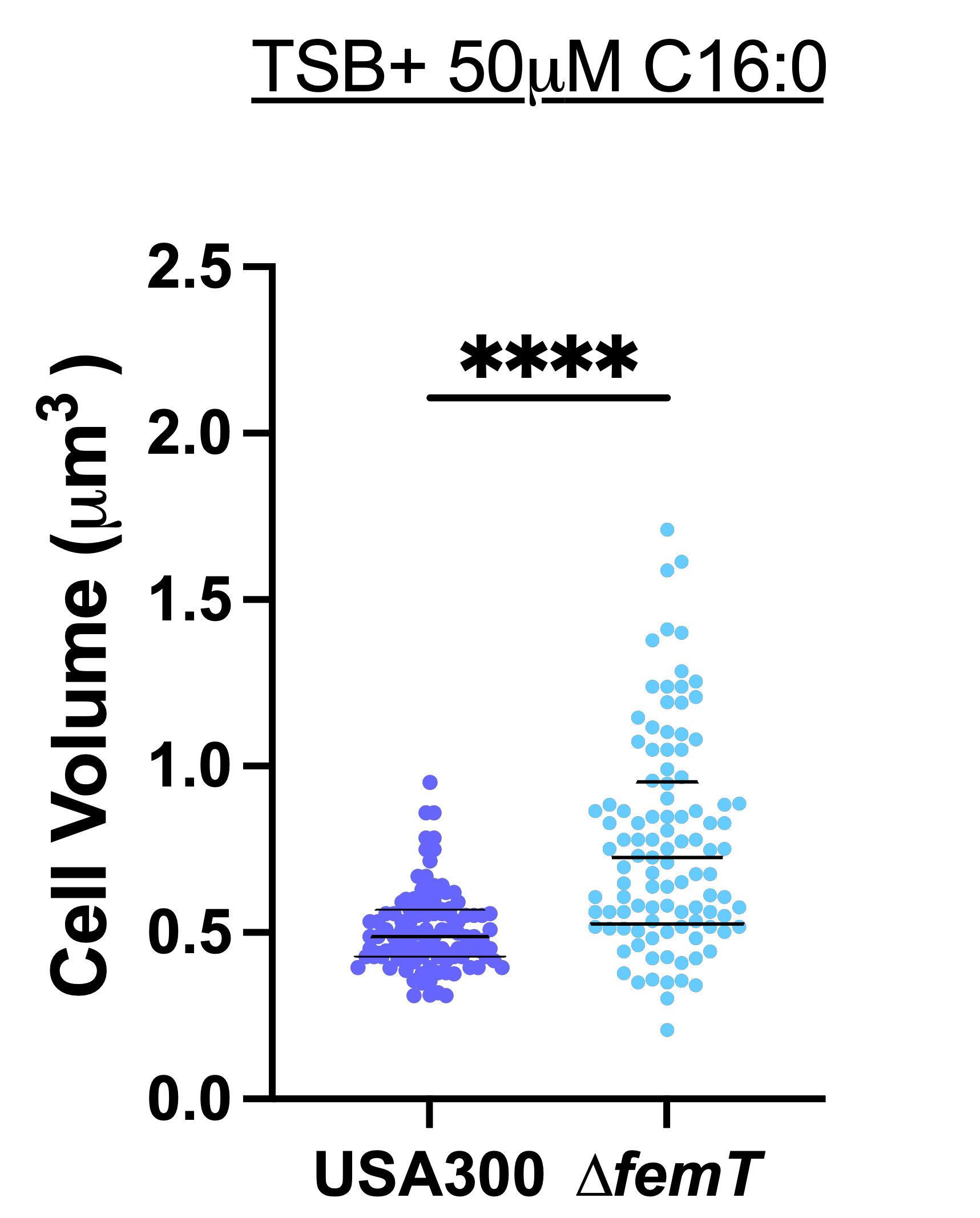

**Supplementary Figure 4. Cell volume measurements of WT and Δ*femT* *Staphylococcus aureus* grown in palmitic acid.** Violin plots with overlaid individual data points represent cell volume distributions measured from 100 cells per strain. Cells were grown in TSB supplemented with 50 µM C16:0 for 3 h, washed and stained with BODIPY-vancomycin. Each dot represents a single cell, with different colors indicating cells analyzed from independent fields of view. Central lines denote the median and interquartile range. Statistical significance between groups was determined using an unpaired two-tailed *t*-test, with significance indicated as **** (p < 0.0001).

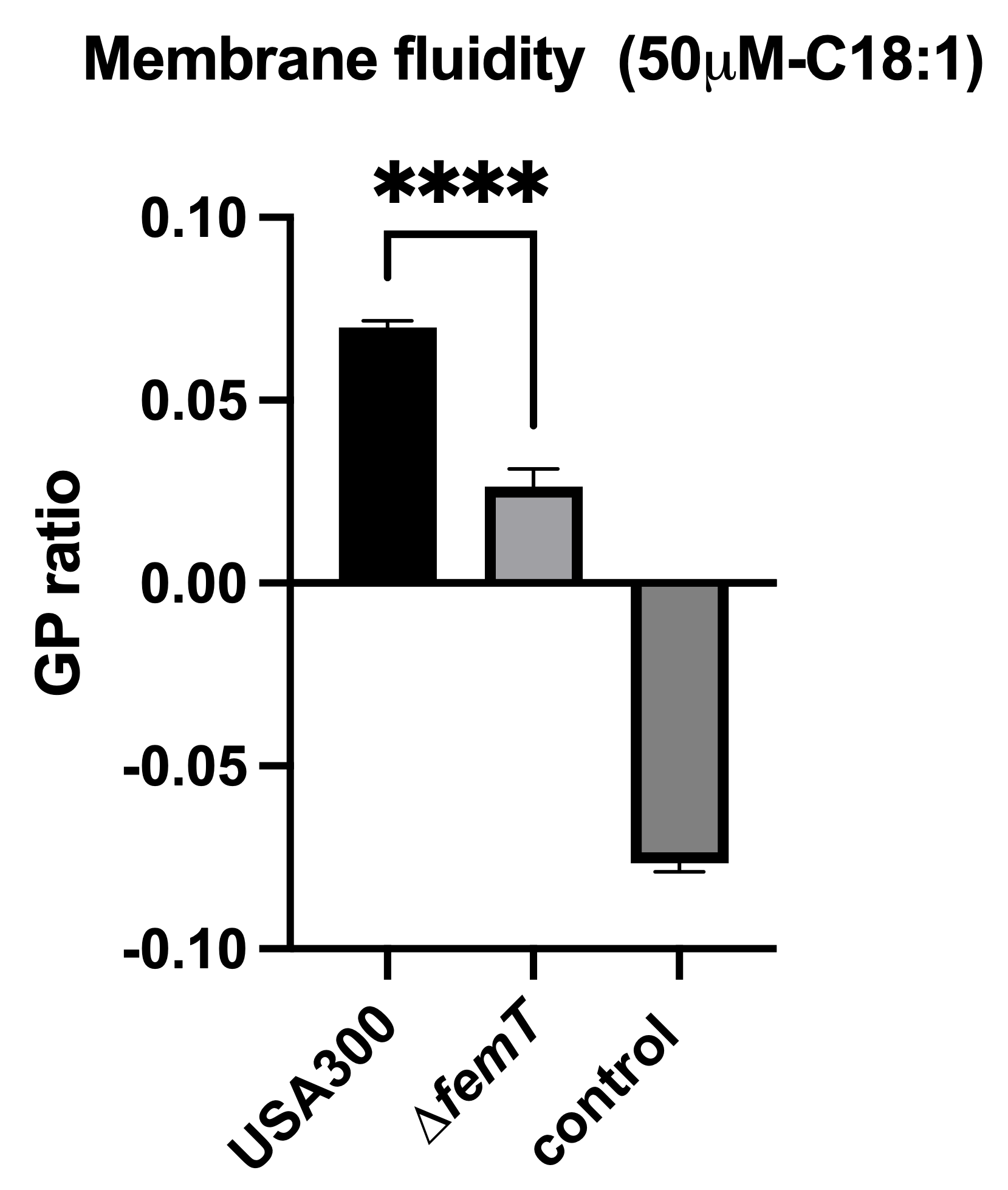

**Supplementary Figure 5: Oleic acid alters membrane fluidity in USA300Δ*femT*.**
Membrane fluidity was assessed by Laurdan generalized polarization (GP) analysis following exposure to 50 µM C18:1. USA300Δ*femT* exhibited a significantly lower GP ratio compared to USA300, indicating increased membrane fluidity in the absence of *femT*. Cells treated with the membrane-fluidizing (50 µM linoleic acid) control displayed strongly reduced GP values. Data represent mean ± SEM from biological replicates. Statistical significance between groups was determined using an unpaired two-tailed *t*-test, with significance indicated as **** (p < 0.0001).

**Table S1. Structural similarity of FemT and FarE with representative transporters determined by DALI.**

| Comparison | Zscore | RMSD (Å) | Aligned residues(lali) | Total residues(nres) | %Sequence identity |
| --- | --- | --- | --- | --- | --- |
| FemT vs SwrC | 60.7 | 3.5 | 1034 | 1065 | 46 |
| FemT vs AdeB | 53.0 | 3.4 | 987 | 1036 | 23 |
| FarE vs MmpL3 | 43.6 | 5.4 | 717 | 944 | 21 |
| Z-score indicates structural similarity; RMSD represents deviation over aligned Cα atoms; lali indicates  aligned residues; nres indicates total residues. Protein structures used were FemT (AF-A0A0H2XER4-F1), SwrC (AF-O31501-F1-v60), AdeB (AF-A0A5N5XY18-F1-v6/ PDB 6OWS), FarE (AF-A0A6B0DA92-F1-v6) and MmpL3 (AF-P9WJV5-F1-v6/ PDB: 7NVH) | | | | | |

**TABLE S2. Strains and plasmids used in this study**

| **Strain / Plasmid** | **Description** | **Source** |
| --- | --- | --- |
| ***S. aureus*** | | |
| RN4220 | Restriction-deficient strain accepting foreign DNA (rk– mk+) | (Novick, 1991) |
| USA300 | CA-MRSA, wild-type strain cured of resistance plasmids | (Arsic *et al.*, 2012) |
| USA300Δ*farE* | Markerless deletion of *farE* | This study |
| USA300Δ*femT* | Markerless deletion of *femT* | This study |
| USA300 pALC2073 | USA300 carrying empty pALC2073 vector; Cmᴿ | This study |
| USA300Δ*femT* pALC2073 | USA300Δ*femT* carrying empty pALC2073 vector; Cmᴿ | This study |
| USA300Δ*femT* pALC*femT* | USA300Δ*femT* complemented with native *femT* on pALC2073; Cmᴿ | This study |
| USA300 pGY*femT::lux* | *femT* promoter fused to *lux*ABCDE reporter; Cmᴿ | This study |
| USA300 pGY*femX::lux* | *femX* promoter fused to *lux*ABCDE reporter; Cmᴿ | This study |
| USA300 pGY*vraX::lux* | *vraX* promoter fused to *lux*ABCDE reporter; Cmᴿ | This study |
| ***E. coli*** | | |
| DH5α | Transformation-competent E. coli cloning strain | Invitrogen |
| BL21(DE3) | Protein expression strain with T7 RNA polymerase | Invitrogen |
| **Plasmid** | | |
| pKOR1 | *E. coli - S. aureus shuttle* vector for allelic replacement; Ampᴿ Cmᴿ Tetᴿ | (Bae and Schneewind, 2006) |
| pKORΔ*femT* | pKOR1 containing ligated PCR products generated with primer pair *femT*UP_*attB1* /*femT*UP_*SacII,* and *femT*DW_*SacII* / *femT*DW_*attB2* | This study |
| pKORΔ*farE* | pKOR1 containing ligated PCR products generated with primer pair *farE*UP_*attB1* /*farE*UP_*SacII* and *farE*DW_*SacII* / *farE*DW_*attB2* | This study |
| pALC2073 | Shuttle vector with xylose/tetO promoter; Cmᴿ Tetᴿ | (Corrigan and Foster, 2009) |
| pALC*femT* | Native *femT* cloned in SacI site; Cmᴿ Tetᴿ | This study |
| pET28a (+) | *E. coli* expression vector for 6×His-tagged proteins; Kmᴿ | Novagen |
| pET:*femT^SD^* | pET28 with soluble exocytoplasmic domains of *femT* cloned in *NheI/SalI*; T7-inducible, Kmᴿ | This study |
| pGY *lux* | Promoterless *lux*ABCDE reporter plasmid; Ampᴿ Cmᴿ | (Mesak, Yim and Davies, 2009) |
| pGY *femX::lux* | *femX* promoter fused to *lux*ABCDE; Ampᴿ Cmᴿ | This study |
| pGY *femT::lux* | *femT* promoter fused to *lux*ABCDE; Ampᴿ Cmᴿ | This study |
| pGY *vraX::lux* | *vraX* promoter fused to *lux*ABCDE; Ampᴿ Cmᴿ | This study |

Abbreviations: Amp^R^ – ampicillin resistance; Cm^R^ – chloramphenicol resistance; Tet^R^ – tetracycline resistance; Km^R^ -kanamycin resistance

**Table S3. Oligonucleotide primers used in this study**

| **Primer** | **Sequenceᵃ** |
| --- | --- |
| *femT*UP_*SacII^b^* | 5′ggacctccgcggCTTATTCCCTAAAGAAAATTGTAATAGC-3′ |
| *femT*UP*_attB1^c^* | 5′attB1-CAAAAGACTACATCCAACACG-3′ |
| *femT*DW_*SacII*^b^ | 5′ggacctccgcggACACTTGTAGTTGACCAGT-3′ |
| *femT*DW_attB2^d^ | 5′attB2-TGTTGCTCACTGTTATCCCTTC-3′ |
| *femX::lux*_F^g^ | 5′TTTcccgggGAGGTGCTATTTATCGCTTTG-3′ |
| *femX::lux*_R^h^ | 5′GTGGCGTTGCACCCGGCATTGTcGacGTAACTGAAATAACTGG-3′ |
| *femT::lux*_F^g^ | 5′GTGGTACAGATAATGATCCcGggAAAGACTCAGAACATTATG-3′ |
| *femT::lux*_R^h^ | 5′GTGGCGTTGCACCCGGCATTGTcGacGTAACTGAAATAACTGG-3′ |
| *vraX::lux_*F^g^ | 5′GTTGGTCCcGgGGATCACGGTGC-3′ |
| *vraX::lux*_Rʰ | 5′GGTGCGCCTTCATGtcGAcACTGTCG-3′ |
| pALC*femT*_Fᵉ | 5′ggtgataaattattttggagctcGTGAAGAAGGGGGAAGTACTGG-3′ |
| pALC*femT*_Rᵉ | 5′cggattacattggagctcCAGTATTGTTTTTAAACACATCG-3′ |
| *femT*-SD1_Fᶦ | 5′GGCGGTGTATATGCtAGcGCTAAATTGAAATTAGAATTACTACC-3′ |
| *femT*-SD1_R | 5′GCTGAAATAAAGCTAGTGCCTAGTCTCGGCTCAACAGGCTTTGCAGTATCCATTG-3′ |
| *femT*-SD2_F | 5′CAATGGATACTGCAAAGCCTGTTGAGCCGAGACTAGGCACTAGCTTTATTTCAGC-3′ |
| *femT*_SD2_Rʰ | 5′-TTTTGtcgacAATCATCTGATGCACCACCGATATT-3′ |

^a^Lowercase nucleotides denote 5′ extensions or nucleotide substitutions introduced during primer design to incorporate restriction enzyme recognition sites. Restriction sequences are underlined.

ᵇ *Sac*II

ᶜ *attB1* site for cloning in pKOR1: GGGGACAAGTTTGTACAAAAAAGCAGGCT

ᵈ attB2 site for cloning in pKOR1: GGGGACCACTTTGTACAAGAAAGCTGGGT

ᵉ*Sac*I

ᶠ*BamH*I

ᵍ*Xma*I

ʰ*Sal*I

^i^*Nhe*I

**REFERENCES**

1. Arsic, B. *et al.* (2012) “Induction of the staphylococcal proteolytic cascade by antimicrobial fatty acids in community acquired methicillin resistant Staphylococcus aureus,” *PloS one*, 7(9).
2. Bae, T. and Schneewind, O. (2006) “Allelic replacement in Staphylococcus aureus with inducible counter-selection,” *Plasmid*, 55(1), pp. 58–63.
3. Corrigan, R.M. and Foster, T.J. (2009) “An improved tetracycline-inducible expression vector for Staphylococcus aureus,” *Plasmid*, 61(2), pp. 126–129. Available at: https://pubmed.ncbi.nlm.nih.gov/18996145/.
4. Mesak, L.R., Yim, G. and Davies, J. (2009) “Improved lux reporters for use in Staphylococcus aureus,” *Plasmid*, 61(3), pp. 182–187.
5. Novick, R.P. (1991) “Genetic systems in staphylococci,” *Methods in enzymology*, 204(C), pp. 587–636.
